# Targeting ALA3 with propiconazole regulates plant growth and enables discovery of promising inhibitor leads

**DOI:** 10.64898/2026.04.17.719095

**Authors:** Qing Gao, Siwei Wang, Dekang Guo, Yunxue Song, Yongsheng Yang, Zhuoyue Chen, Xianyu Zhang, Ronghua Chen, Hanhong Xu, Fei Lin

## Abstract

Propiconazole (PCZ) is widely misused growth regulator in leafy *Brassica* vegetables. Developing green strategies for managing plant architecture has become an urgent agricultural priority. Here, we identified from a membrane-protein-defective yeast library a P4-ATP phospholipid flippase, aminophospholipid ATPase 3 (ALA3), as a target sensitive to PCZ. ALA3 exhibits high binding affinity for PCZ, which inhibits its ATPase activity. Knockdown of *ALA3* rendered yeast, *Arabidopsis*, and *Brassica rapa* less sensitive to PCZ and conferred a growth-inhibited phenotype. This dwarfing phenotype is mediated through the interaction between ALA3 and CYP51G1 that jointly acts within the brassinosteroid regulatory pathway. Furthermore, we identified lead compounds A01 and A15 as ALA3-targeting agents, and compared to PCZ, they display superior binding affinity and reduced toxicity. Our work establishes ALA3 as a key mediator of PCZ-induced dwarfism and provides dual strategies—creating promising varieties through gene editing and developing targeted green pesticides—to reduce PCZ use.

**Teaser:** Targeting ALA3 reduces PCZ use through gene-edited varieties and green pesticides.

## Introduction

In modern agriculture, the frequency of plant growth regulator applications has risen steadily to meet market demand for desirable leafy vegetables. This trend is particularly concerning given the intrinsic characteristics of leafy vegetables, such as their short growth cycles, high dietary consumption frequency, susceptibility to pests/diseases, and large pesticide-exposed surface areas (*1–3*). These traits collectively heighten the risks associated with pesticide exposure and safety. Agricultural practice has demonstrated that propiconazole (PCZ) application provides dual benefits in leafy *Brassica* vegetables cultivation: broad-spectrum fungicidal activity and growth modulation that enhances marketability by dwarfing and suppressing premature flowering in plants. PCZ has been adopted by farmers to promote compact plant architecture in leafy *Brassica* vegetables, achieving growth restraint, stem thickening, and pigmentation enhancement (*4–6*). The application of PCZ often fails to observe a sufficient safety interval, leading to its accumulation and residue in crops and thereby posing a threat to dietary safety (*4, 5, 7*). Consequently, establishing green and sustainable strategies for plant growth modulation has become urgent agricultural priorities.

At present, clarifying the molecular mechanism through which PCZ regulates the plant architecture of leafy *Brassica* vegetables is key to addressing this production challenge and constitutes the essential first step toward establishing green and safe growth-regulation strategies. The fungicidal action of PCZ, a triazole fungicide, has been characterized, it primarily inhibits fungal growth by targeting CYP51 (Sterol 14α-demethylase) to block ergosterol biosynthesis and disrupt membrane integrity (*8, 9*). Sterol 14α-demethylase, the most evolutionarily conserved cytochrome P450 family, catalyzing the first step following cyclization in sterol biosynthesis and ultimately leading to the formation of brassinosteroids (BRs) in plants (*10*). Rescue experiments with BR biosynthetic intermediates coupled with *in vitro* protein inhibition assays demonstrated that PCZ specifically suppresses C22/C23 side-chain hydroxylation during campesterol-to-teasterone conversion (*11*). Furthermore, it has been reported that BR rescue the phenotypic defects induced by PCZ, leading to the proposal that PCZ acts as a specific inhibitor of BR (*12*). While the role of PCZ in regulating plant growth through modulation of phytosterol and BR biosynthesis has been documented, the molecular mechanisms by which PCZ enters cells and induces membrane structural alterations remain unclear. Therefore, as an exogenous compound, the cellular uptake of PCZ and its subsequent trafficking to the smooth endoplasmic reticulum (ER) and mitochondrial inner membrane to interact with cytochrome P450 enzymes require further mechanistic elucidation. Previous studies have documented PCZ-induced alterations in membrane architecture. In *Tradescantia virginiana*, PCZ inhibits pollen germination and tube elongation while disrupting cytoplasmic streaming and cytoskeletal organization, suggesting its effects may be mediated through perturbations in membrane lipid composition or integrity that indirectly impair cytoskeleton-membrane interactions rather than through direct targeting of microfilaments or microtubules (*13*). Additionally, given that the primary target of BRs is the plasma membrane (PM)-localized receptor complex BRI1/BAK1 (*14*), it suggests that PCZ may exert its effects by influencing membrane integrity or interfering with PM-initiated signal transduction. Collectively, these findings point to the PM as a responsive site for PCZ, initiating a series of downstream cellular events.

P4-ATPase phospholipid flippases are key regulators for maintaining membrane functionality, and their dysfunction has been associated to disease (*15*). In contrast, their interactions with small molecules have rarely been reported. A pivotal breakthrough came in 1996 with the identification of yeast P4-ATPase Drs2p, whose deletion impaired phosphatidylserine (PS) translocation from the exoplasmic to cytoplasmic leaflet (*16*). Subsequent studies revealed that Drs2p forms a functional complex with its *β*-subunit Cdc50p at the Golgi-trans Golgi network (TGN), where it orchestrates clathrin-coated vesicle biogenesis through maintenance of membrane phospholipid asymmetry (*17, 18*), demonstrating that P4-ATPases serve not merely as lipid “transporters” but as fundamental “architects” of membrane trafficking systems. In 2019, cryo-EM structures of yeast Drs2p-Cdc50p were resolved at resolutions of 2.8 Å, 3.7 Å, and 2.9 Å in the E2P^inhib^, E2P^inter^, and E2P^active^ conformations, respectively (*19, 20*), these structures elucidated the mechanisms of autoinhibition and phosphatidylinositol 4-phosphate (PI4P)-dependent activation, proposing a self-regulatory model for lipid entry (*20*).

The plant *ALA3* gene exhibits multifaceted membrane-associated regulatory properties. As a homolog of DRS2 (*21*), ALA3 localizes to the PM, TGN, and Golgi (*22*). *Arabidopsis thaliana* (*A. thaliana*) *ala3* mutants display compact rosette leaves with rounded morphology and impaired root and hypocotyl growth(*21*). The *ALA3* in *Arabidopsis* specifically drives secretory vesicle generation in root cap peripheral columella cells (*21*). *ALA3* also facilitates the maintenance of auxin gradients and regulates polar transport cycles, thereby ensuring normal lateral root development and pollen tube growth (*22, 23*). Notably, the potential of this gene in pesticide science and broader chemical applications remains largely unexploited. Thus, our study provides an innovative bridge between molecular biology and chemical sciences.

Addressing the critical agricultural need for controlled-growth and safe production of leafy *Brassica* vegetables, this study elucidates the molecular mechanism by which PCZ modulates plant architecture through targeting the membrane protein ALA3. Our findings demonstrate that PCZ treatment inhibits excessive vegetative growth, identifying ALA3 as a sensitive membrane target for PCZ. PCZ exhibits strong binding affinity to ALA3, thereby suppressing ATPase activity. ALA3 interacts with CYP51G1 and ALA3 may be involved in BR biosynthetic pathway. Lead compounds targeting the binding pocket between PCZ and ALA3 expected to serve as the basic framework for designing new pesticides. These discoveries establish dual strategies for growth modulation: ALA3-targeted gene editing and structure-based pesticide development, offering sustainable alternatives for precision agriculture.

## Results

### PCZ suppresses the growth of *A. thaliana*

To investigate whether *A. thaliana* develops compact-leaf phenotypes analogous to those observed in field-grown leafy *Brassica* vegetables following PCZ treatment, we carried out a pot experiment. Phenotypic analysis of *A. thaliana* revealed severe growth retardation, characterized by a compact architecture with markedly suppressed expansion of newly emerged stems and leaves following PCZ treatment (fig. S1A). Mature leaves exhibited significantly shortened petioles and reduced leaf length (fig. S1, B and C), while leaf width remained unaltered (fig. S1D). The total length of leaf was substantially diminished (fig. S1E). These collective findings demonstrate PCZ can effectively control excessive vegetative growth while promoting compact plant architecture.

### Screening of yeast mutant library identified ALA3 as PCZ-sensitive protein

A yeast mutant library of membrane transporters was assembled to screen for PCZ-associated proteins. The result showed that *drs2Δ* mutant (lacking the aminophospholipid-translocating P4-type ATPase DRS2) (*19*) exhibited marked resistance as PCZ concentration increased, while other deletion mutants showed growth inhibition (fig. S2). Phylogenetic and homologous comparative analyses identified *A. thaliana* AtALA3 (AT1G59820) as homolog of yeast DRS2 (YAL026C)(*21*), with 42.31% amino acid sequence identity (fig. S3 and table S1).

Expression of *AtALA3* in *drs2Δ* and BY4741 yeast strains altered their sensitivity to PCZ (*24–26*), revealing that yeast strains expressing *AtALA3* exhibited significantly impaired growth on PCZ-containing media compared to pYES2-empty vector controls, with overexpression strains showing more pronounced growth suppression (Fig. 1A). Parallel liquid culture assays monitoring OD_600_ revealed identical trends, with *AtALA3*-expressing strains displaying markedly reduced proliferation rates versus three control groups (Fig. 1, B and C). These collective data demonstrate that *DRS2* deficiency confers PCZ resistance in yeast, heterologous expression of plant *AtALA3* restores PCZ sensitivity, *AtALA3* overexpression enhances susceptibility. Thus, ALA3 is conclusively identified as a PCZ-sensitivity determinant.

**Fig. 1.**
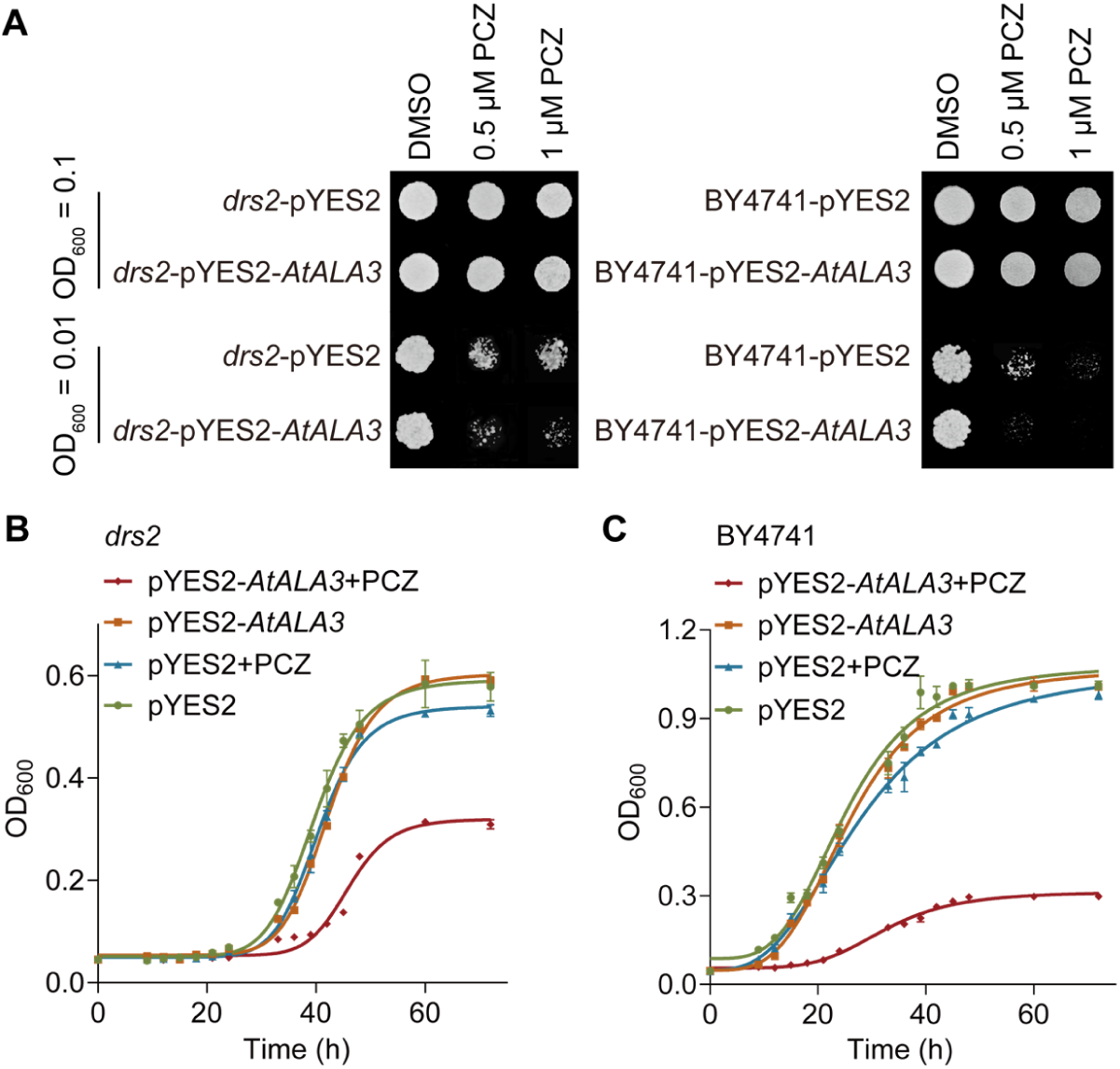
Identification of sensitivity of *AtALA3* heterologous expression yeast strain to propiconazole (PCZ). (**A**) Growth of *drs2* and BY4741 yeast strains supplemented homologous gene *AtALA3* on SD/-His-Leu-Met medium containing DMSO, 0.5 μM PCZ, or 1.0 μM PCZ. Plates were photographed after 3 days of incubation at 30°C. (**B** and **C**) Growth curve of *drs2* (B) and BY4741 (C) yeast strains supplemented homologous gene *AtALA3* in SD/-His-Leu-Met liquid medium containing DMSO or 0.5 μM PCZ at 30°C. The growth of yeast strains was monitored by OD_600_ readings over 72 hours. In (B) and (C), the error bars indicate the SD (*n* = 3). BY4741 is the wild-type yeast strain; *drs2* is a mutant in the BY4741 background.

### PCZ docks into and likely occupies the PI4P binding pocket of ALA3

To elucidate the mechanistic basis of the interaction between PCZ and ALA3, we generated a homology model of the AtALA3-AtALIS1 complex based on the DRS2-Cdc50p template (*20*) and performed molecular docking with PCZ. The ALIS1-ALIS5 protein family (ALA3’s *β*-subunits) represents functional equivalents of yeast Cdc50p (*21*). Based on reported studies, the final localization and lipid substrate specificity of P4-ATPases are independent of the properties of their cognate ALIS *β*-subunits that mediate these interactions (*27*), we selected *A. thaliana* AtALIS1 as the *β*-subunits for modeling. Molecular docking of PCZ into the AtALA3-AtALIS1 model revealed that PCZ binds within a hydrophobic pocket formed by transmembrane helices TM7, TM8, and TM10, as well as the autoinhibitory C-terminus. The binding energy was calculated to be −8.2 kcal/mol, with a ligand efficiency of 0.3737 kcal/(molꞏK). PCZ engages in non-covalent interactions with surrounding residues: the N atom of its triazole ring forms a hydrogen bond with Asparagine 1044 (Asn-1044), the dioxolane group forms hydrophobic-interaction with Phenylalanine 1112 (Phe-1112) and Glutamine 1113 (Gln-1113) in TM10, and the para-substituted phenyl ring and propyl side chain’s chlorine atom form a halogen bond with Tyrosine 1124 (Tyr-1124) in the autoinhibitory C-terminus (Fig. 2A). Molecular docking of AtALA3-AtALIS1 complex with PCZ predicted critical binding residues in the active pocket, which were subsequently validated by site-directed mutagenesis (table S3). The point-mutant showed reduced yeast growth sensitivity to PCZ compared to strain expressing *AtALA3* (Fig. 2B), confirming the essentiality of these residues for target interaction.

**Fig. 2.**
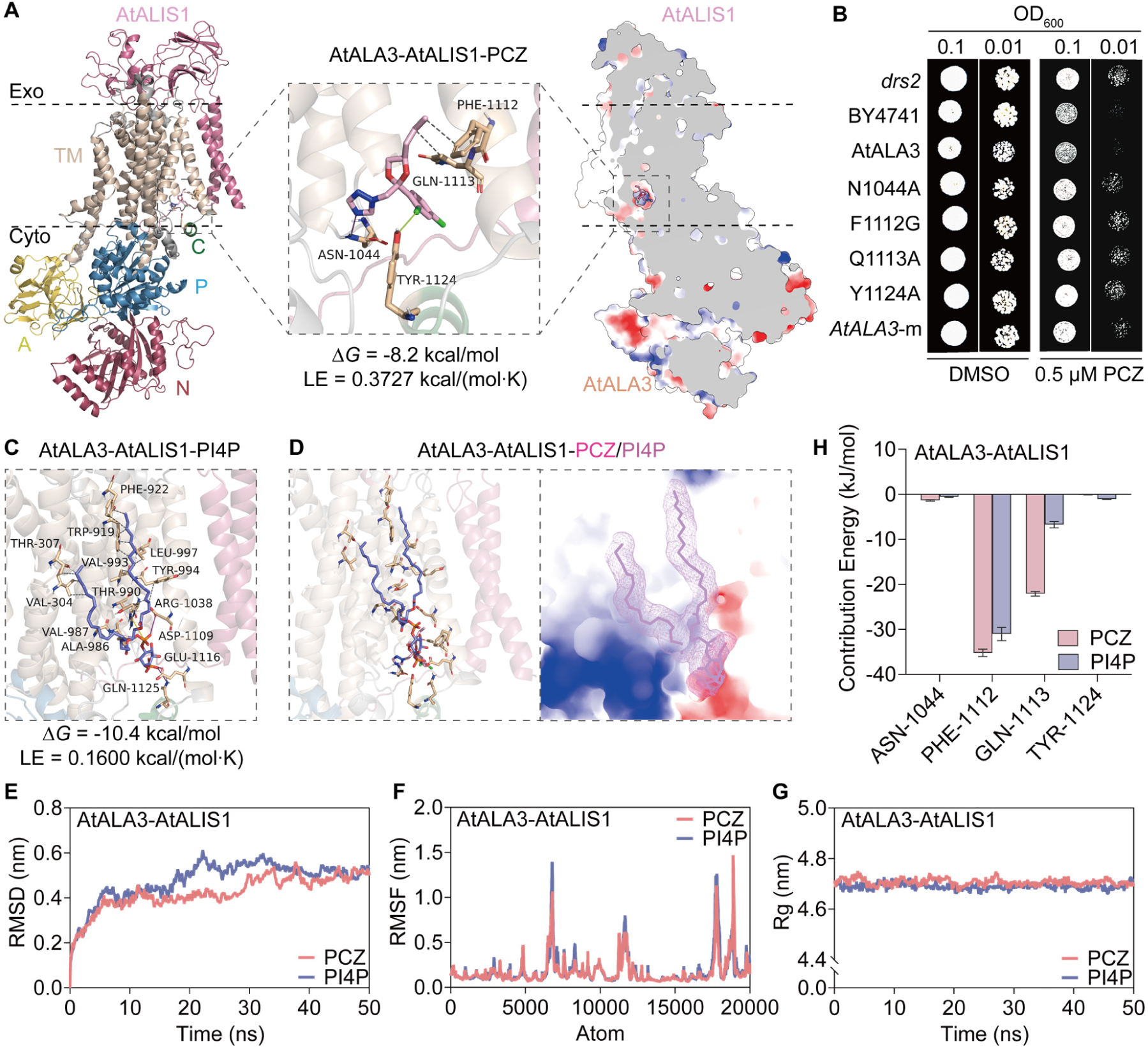
Molecular docking, interaction site validation, and molecular dynamics simulation analysis of AtALA3-AtALIS1 protein binding to PCZ and PI4P. (**A**) Molecular docking between AtALA3-AtALIS1 and PCZ. From left to right are cartoon diagrams of molecular docking, binding site visualization and surface cross-sections. (**B**) Growth of *drs2* yeast strains supplemented *AtALA3* and site-specific mutation type on SD/-His-Leu-Met medium containing DMSO or 0.5 μM PCZ. Plates were photographed after 3 days of incubation at 30°C. Mutant designations indicate yeast expressing *AtALA3* with the specified amino acid substitution: N1044A (Asn1044Ala); F1112G (Phe1112Gly); Q1113A (Gln1113Ala); Y1124A (Tyr1124Ala). “atala3-m” denotes the strain containing all four mutations. (**C**) Molecular docking between AtALA3-AtALIS1 and PI4P. (**D**) Schematic representation of spatial structure of PCZ and PI4P robbing AtALA3-AtALIS1 pockets. (**E**) The Root Mean Square Deviation (RMSD) curves of AtALA3-AtALIS1 protein and ligand (PCZ/PI4P) complexes over time. (**F**) The Root Mean Square Fluctuation (RMSF) curves of protein skeleton atoms in AtALA3-AtALIS1 protein and ligand (PCZ/PI4P) complexes. (**G**) The Radius gyration (Rg) of AtALA3-AtALIS1 protein and ligand (PCZ/PI4P) complexes over time. (**H**) Free energy contribution of key amino acids residues in AtALA3-AtALIS1-PCZ/PI4P protein-substrate complex. In (A) and (C), △*G* represents binding energy. LE represents ligand efficiency. PCZ (pink), and PI4P (purple) are represented by the stick model. Gray dashed lines represent hydrophobic-Interactions, purple solid lines represent hydrogen-bonds, yellow dashed lines represent π-stacking, green solid lines represent halogen-bonds, orange dashed lines represent π-cation-interactions, yellow solid lines represent salt-bridges. In (H), the error bars indicate the SD.

P4-ATPase functionality requires PI4P-dependent activation (*28*) to drive the transbilayer flipping of specific phospholipid substrates such as PS from the exoplasmic to the cytoplasmic leaflet (*20*). Following homology modeling of the membrane protein complexes AtALA3, we performed molecular docking of PI4P with the ALA3 dimeric proteins. The results revealed that PI4P binds to the active site of ALA3, formed by transmembrane helices TM7, TM8, TM10, and the autoinhibitory C-terminal hydrophobic pocket. The binding energy was calculated to be −10.4 kcal/mol, with a ligand efficiency of 0.1600 kcal/(molꞏK) (Fig. 2C). Visualization of the co-docked conformation demonstrated that PCZ occupies the same binding pocket in ALA3 that is normally utilized by PI4P (Fig. 2D). Although the binding energy of PCZ to ALA3 was marginally lower than that of PI4P, its ligand efficiency significantly surpassed that of PI4P (Fig. 2, A and C).

Using root mean square deviation (RMSD), root mean square fluctuation (RMSF), and radius gyration (Rg) from molecular dynamics (MD) simulations to assess the conformational stability of the docked complex, we observed a major conformational shift between 20 ns and 30 ns. Thereafter, the simulation curves plateaued, indicating structural convergence into a stable conformational state (Fig. 2, E to G). Moreover, the dynamic fluctuations of PCZ and PI4P were highly similar, supporting the hypothesis that the two molecules compete for binding, as proposed above (Fig. 2, E to G). Free energy decomposition analysis identified key residues with binding energy contributions (≤ −10 kcal/mol) for both PCZ and PI4P complexes: Phe-1112 and Gln-1113 in AtALA3-AtALIS1. Notably, these residues exhibited stronger binding energy contributions for PCZ than for PI4P (Fig. 2H), indicating that PCZ likely competes with PI4P for binding to these critical sites within the active pocket, where these key residues stabilize the complexes primarily through non-covalent interactions, supporting a competitive occupation mechanism. Collectively, these findings suggest that exogenously applied PCZ may occupy the same binding pocket in ALA3 that is normally utilized by the lipid flippase regulator PI4P, which could thereby impede the activation of its autoinhibitory conformational mechanism.

### PCZ interacts with ALA3 and inhibits its ATPase activity

To validate the interaction predicted by molecular docking simulations and to determine whether PCZ activates or inhibits ALA3 function, we performed interaction assays at protein levels. Membrane protein expression using the BJ5465 yeast system (fig. S5) and subsequent SDS-PAGE densitometry indicated decreased protein levels following PCZ treatment (fig. S6A). Surface plasmon resonance (SPR) analysis revealed the binding affinities between PCZ and ALA3 proteins. Concentration-dependent SPR binding signals (10 nM to 2560 nM) demonstrated enhanced PCZ interactions with ALA3 homologous proteins, generating characteristic SPR binding kinetics curves (Fig. 3, A and B), no binding signals were detected in control experiments (fig. S6, B to D). The equilibrium dissociation constants (Avg KD) for DRS2, AtALA3 were 3.09 × 10^-8^ M and 2.04 × 10^-8^ M with PCZ respectively, indicating strong binding affinities (Fig. 3C). These results demonstrate significant binding affinities between ALA3 membrane proteins and PCZ. *In vitro* ATPase activity was assayed to validate the bioactivity of the obtained protein complex. In the presence or absence of phosphatidylserine (POPS), the phosphate released from ATP hydrolysis was detected colorimetrically. The ATPase activity of the protein complex under different conditions was quantified by calculation. The results show that PCZ inhibits ALA3 enzymatic activity, with enhanced suppression observed in the presence of the phospholipid POPS (Fig. 3, D and E). Collectively, these findings demonstrate that PCZ exhibits high binding affinity for ALA3 and suppresses its activity.

**Fig. 3.**
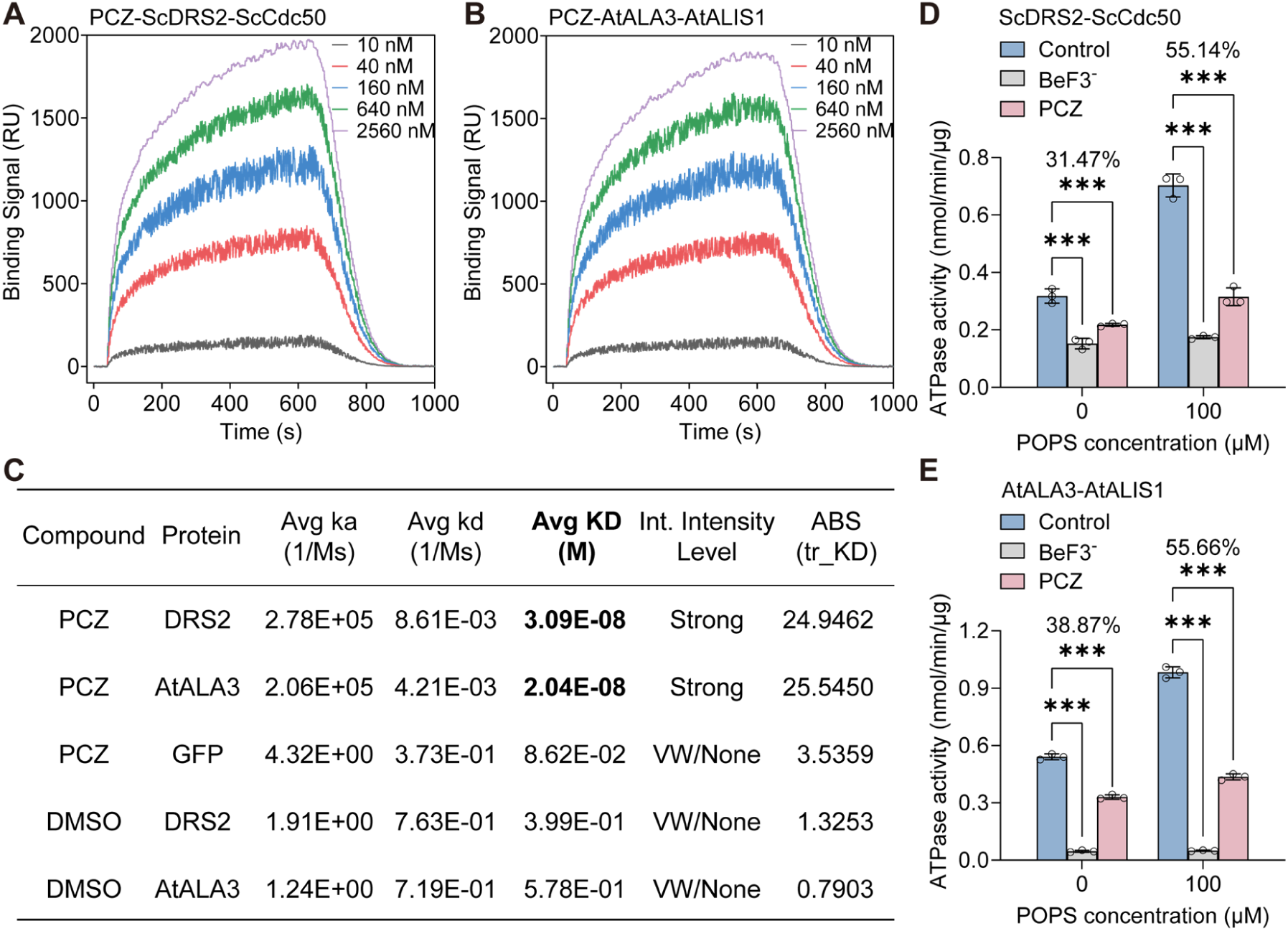
Validation of PCZ-ALA3 binding affinity and PCZ-mediated ATPase activity inhibition. (**A** and **B**) Surface plasmon resonance (SPR) affinity binding signals between PCZ and ScDRS2-ScCdc50 (A), AtALA3-AtALIS1 (B). (**C**) SPR affinity coefficients of PCZ and ScDRS2-ScCdc50, AtALA3-AtALIS1. Avg Ka (1/Ms): The average association rate constant (Ka) represents the ratio of complex formation per unit time relative to initial reactant concentrations, where higher values indicate faster molecular binding. Avg Kd (1/s): The mean dissociation rate constant (Kd) reflects the proportion of complex dissociation per unit time, with elevated values corresponding to faster complex disintegration. Avg KD (M): The equilibrium dissociation constant (KD = Kd/Ka) quantifies binding affinity at dynamic equilibrium. Lower KD values signify stronger intermolecular interactions. Interaction Intensity Level: Determination of affinity. KD range: 10^-^¹³-10^-^⁵ M = strong; 10^-^⁵-10^-^³ M = moderate; 10^-^³-2×10^-^² M = weak; >2×10^-^² M = negligible. ABS (tr_KD): Absolute affinity coefficient. ABS (tr_KD) = ABS (log_2_KD), where higher numerical values indicate enhanced binding affinity. Reported as mean ± SD from four technical replicates. (**D** and **E**) Functional validation of P4-ATPase protein complex enzymatic activity. Phosphatidylserine (POPS) is a substrate for ALA3’s phospholipid flippase activity. BeF3^-^, a known inhibitor of P-type ATPases, was used as a negative control. In (D) and (E), the error bars indicate the SD (*n* = 3). Asterisks indicate significant differences between mean values (two-way ANOVA: Dunnett, \**P* < 0.05, \*\**P* < 0.01 and \*\*\**P* < 0.001).

### *ALA3* loss-of-function mutants in *A. thaliana* exhibit reduced sensitivity to PCZ

To evaluate PCZ sensitivity phenotypes, we selected distinct T-DNA insertion mutants (fig. S7A), including *atala3-p*, *atala3-l* (fig. S7B), and *atala3-4* mutant (*21, 23*). All mutants displayed more compact architecture than Col-0 (Fig. 4A), with significantly reduced petiole length, leaf length, and leaf width (Fig. 4, B to D). Relative to PCZ-treated wild-type plants, *atala3-p* showed similar growth inhibition, while *atala3-4* exhibited more severe growth defects, and *atala3-l* displayed distinct petiole length and leaf width phenotypes (Fig. 4, A to D). To address the practical need for viable seeds in crop production, we assessed the fertility of the *Arabidopsis atala3* mutant, even though our core research targets vegetative growth regulation. The mutants produced shorter siliques but was fully fertile and yielded seeds (fig. S7, C and D). The expression analysis of *AtALA3* in the mutants showed that the transcript level in *atala3-p*, which harbors a T-DNA insertion in the promoter, was the lowest compared to *atala3-l* (insertion in the last exon) and *atala3-4* (insertion in the second intron) (fig. S7E). This mutation does not disrupt the full-length protein structure but drastically reduces mRNA abundance, confirming *atala3-p* as an effective gene-silencing mutant caused by promoter insertion. Phenotypic analysis indicated that promoter-mediated expression modulation has subtler effects, leading to our selection of *atala3-p* (hereafter *atala3*) as the primary low-expression line. The results of 50 mg/L PCZ treatment on 4-week-old *A. thaliana* showed that, in wild-type adult plants, PCZ treatment significantly shortened the petioles and reduced both leaf length and width. The *atala3* mutant itself also exhibited shorter petioles and significantly reduced leaf length and width compared to the wild type. When the mutant was treated with PCZ, petiole length and leaf width showed a slight, non-significant decrease, while leaf length was still significantly reduced—though the inhibition rate was lower than that in PCZ-treated wild-type plants (fig. S8A, C to E). Furthermore, PCZ treatment and *atala3* mutant delayed the date of bolting. By day 9 post-bolting, bolt height was significantly reduced across all treated groups. Notably, the magnitude of height suppression was attenuated in PCZ-treated *atala3* mutants compared to treated wild-type plants (Fig. 4E). These findings indicate that disrupting *ALA3* reduces plant sensitivity to PCZ.

**Fig. 4.**
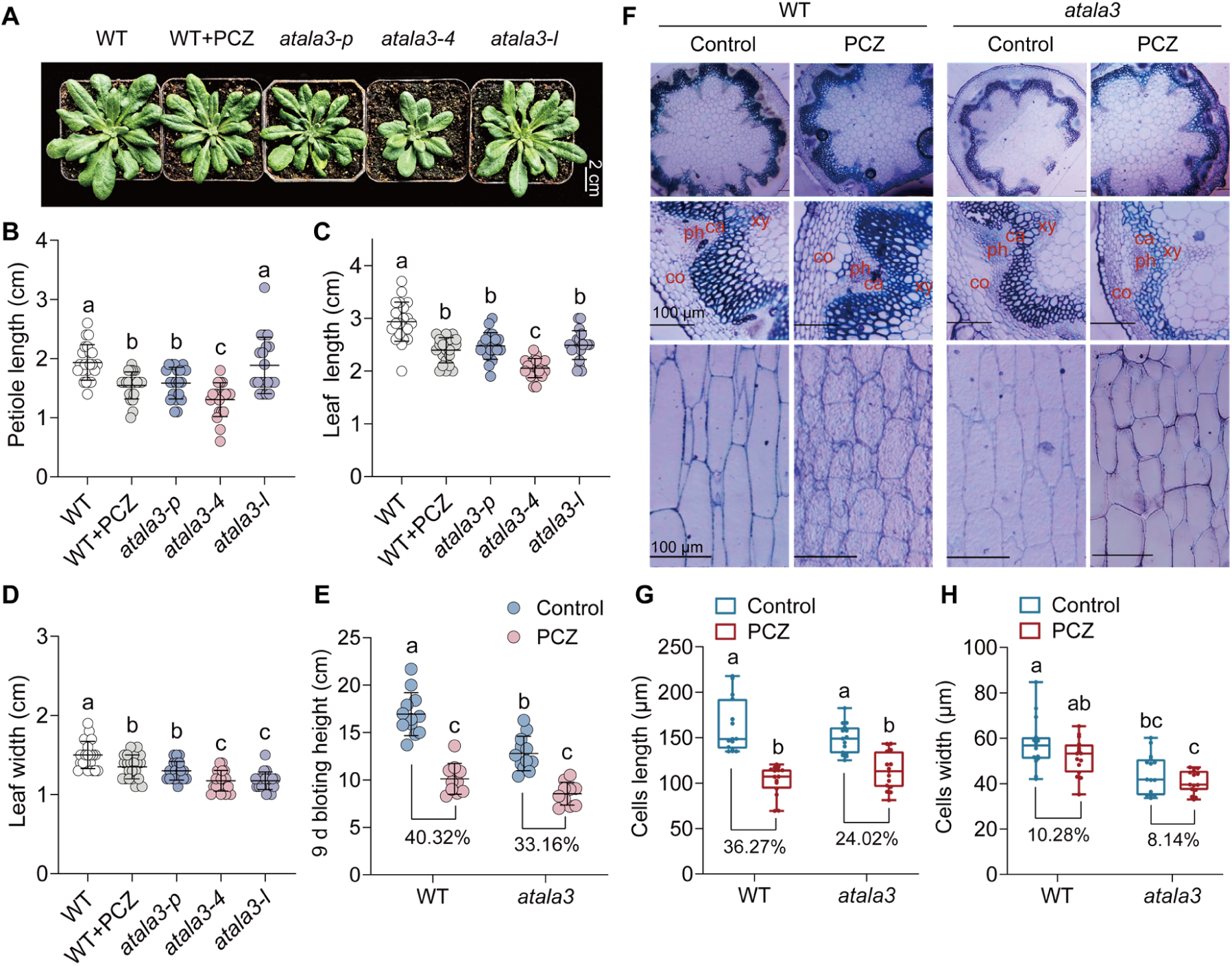
Phenotypic and microstructural analysis of *atala3* mutant. (**A**) Phenotypes of WT (Col-0) and *atala3-p/4/l* mutant (28-day-old). Scale bar, 2 cm. (**B** to **D**) Comparison of mature leaf petiole length (B), leaf length (C), and leaf width (D) with or without 50 mg/L PCZ-treated WT and *atala3-p/4/l* mutant (28-day-old). (**E**) Comparison of 9 d bolting height of control-treated or 50 mg/L PCZ-treated WT and *atala3-p* mutant (28-day-old). (**F**) Resin slice cross sections and longitudinal sections of control-treated or PCZ-treated WT (Col-0) and *atala3* (*atala3-p*) mutant stem base. Scale bar, 100 μm. xy-xylem, ca-cambium, ph-phloem, co-cortex. (**G** and **H**) Comparison of parenchymatous cells length (G) and width (H) of control-treated or PCZ-treated WT and *atala3* mutant stem base. In (B) to (D), the error bars indicate the SD (*n* ≥ 20). In (E), the error bars indicate the SD (*n* ≥ 10). In (G) and (H), the whiskers indicate min to max (*n* = 15). Letters indicate significant differences between mean values (one-way ANOVA: Tukey, *P* < 0.05). Percentages represent the decrease or increase of each indicator.

Histological sections showed reduced xylem and increased phloem in both PCZ-treated Col-0 and untreated *atala3* stems, with mutant phloem remaining unchanged after PCZ treatment (Fig. 4F). Longitudinal sections revealed compacted parenchyma cells in all PCZ-treated plants (Fig. 4F), cell length and width are both inhibited, but the mutant has a lower inhibition rate than Col-0 (Fig. 4, G and H), confirming its reduced PCZ sensitivity. In summary, the phenotypic and microstructural changes of *atala3* mutant indicate that reducing ALA3 expression decreases sensitivity to PCZ in *A. thaliana*, which is consistent with the conclusion of yeast heterologous expression (Fig. 1).

### ALA3 interacts with sterol 14α-demethylase CYP51G1

Early studies established that PCZ inhibits fungal growth by targeting sterol 14α-demethylase (CYP51), thereby blocking ergosterol biosynthesis and compromising membrane integrity (*8, 9*). To further dissect the mechanism by which PCZ regulates plant growth, the interaction between ALA3 and the plant sterol 14α-demethylase CYP51G1 was first assessed using the yeast split-ubiquitin system. The result of yeast two hybrid (Y2H) showed that only yeast co-expressing AtALA3 and AtCYP51G1 grew and turned blue on both selective media, while control co-transformants with empty vectors failed to grow (Fig. 5A), demonstrating a specific physical interaction between AtALA3 and AtCYP51G1.

**Fig. 5.**
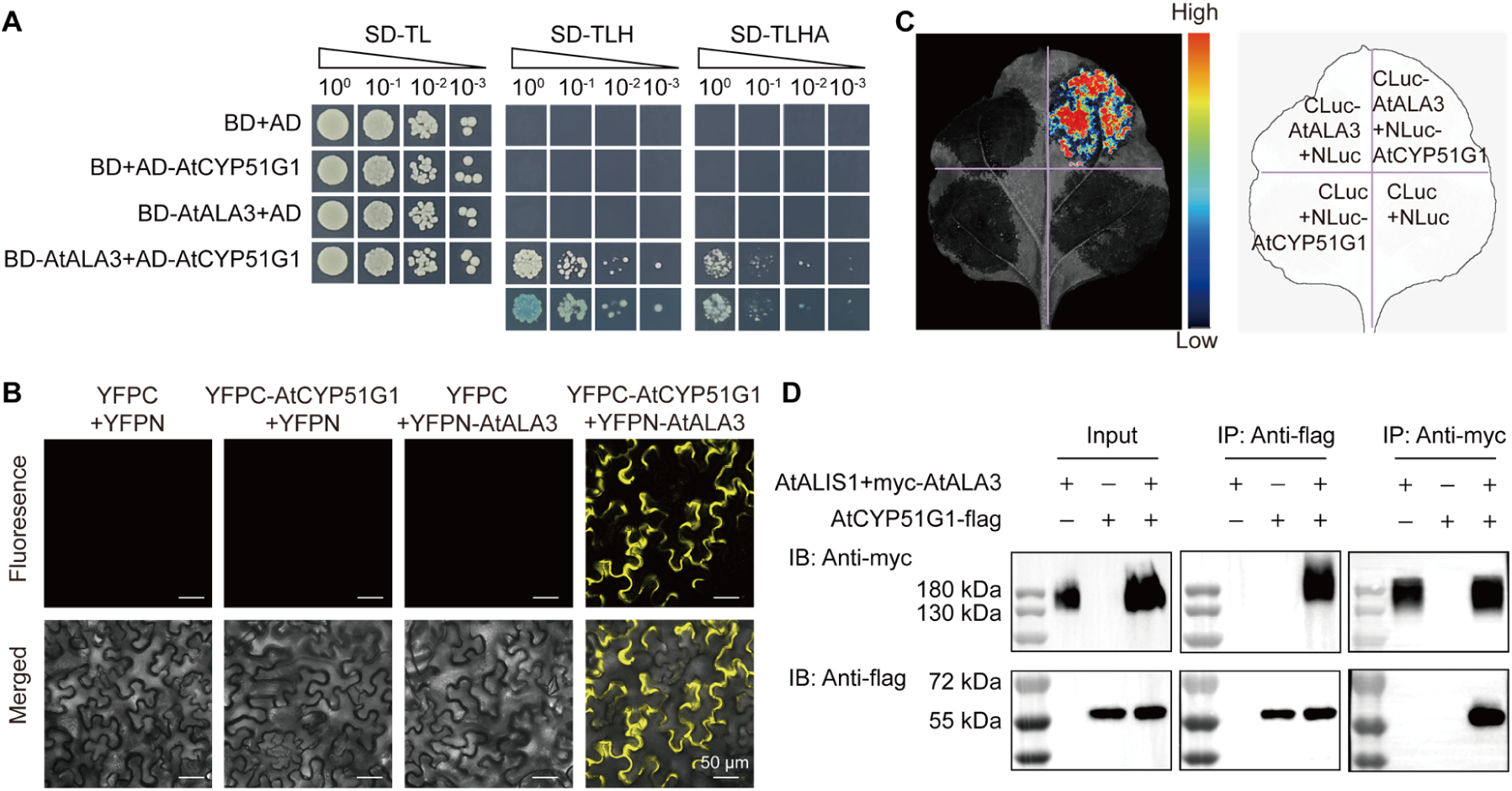
The Interaction between AtALA3 and AtCYP51G1. (**A**) AtALA3 interacts with AtCYP51G1 in yeast two-hybrid (Y2H) system. The 1:10 serial dilutions of yeast cells were spotted on control medium (SD-TL) or selective medium (SD-TLH and SD-TLHA). The same set of yeast transformants were assayed for X-gal activities. SD-TL: SD/-Trp-Leu, SD-TLH: SD/-Trp-Leu-His+30 mM 3AT, SD-TLHA: SD/-Trp-Leu-His-Ade+30 mM 3AT. AD: Activation Domain vector, BD: Binding Domain vector. (**B**) *N. benthamiana* leaves coexpressing AtALA3-YFPN and AtCYP51G1-YFPC were examined by confocal laser scanning microscopy, excitation wavelength is 514 nm. Empty vector-transfected samples are negative controls. Scale bar, 50 μm. (**C**) *N. benthamiana* leaves coexpressing CLuc-AtALA3 and NLuc-AtCYP51G1 were examined by catalytic imaging using potassium fluorescein as substrate. Empty vector-transfected sections are negative controls. The legend indicates fluorescence intensity values. (**D**) Co-IP analysis for AtALA3 and AtCYP51G1 interaction in agrobacterium-infiltrated *N. benthamiana* leaves. myc-AtALA3 and AtCYP51G1-flag were coexpressed as the indicated combinations (Top). “-” represents vector alone. Input (left) are controls. Interaction proteins (middle) were immunoprecipitated with anti-flag antibody (IP: flag) and detected with anti-myc antibody (IB: myc). Also, interaction proteins (right) were immunoprecipitated with anti-myc antibody (IP: myc) and detected with anti-flag antibody (IB: flag). In (B-D), different *Agrobacterium* combinations from the same experiment were infiltrated at adjacent locations on a single *N. benthamiana* leaf.

To confirm the interaction between AtALA3 and AtCYP51G1, we performed a bimolecular fluorescence complementation (BiFC) assay in *Nicotiana benthamiana* (*N. benthamiana*) leaves. Confocal microscopy revealed strong YFP fluorescence in leaves co-expressing AtALA3 and AtCYP51G1, whereas no signal was detected in controls co-infiltrated with empty vectors (Fig. 5B), indicating a specific interaction between AtALA3 and AtCYP51G1 that facilitates YFP reassembly.

The interaction between AtALA3 and AtCYP51G1 was further verified using a luciferase complementation assay (LCA). The result of LCA revealed that luminescence was detected in leaves co-expressing AtALA3 and AtCYP51G1, but not in control leaves co-infiltrated with empty vectors (Fig. 5C), demonstrating a specific interaction between AtALA3 and AtCYP51G1.

Co-immunoprecipitation (Co-IP) assays were performed to examine the interaction between AtALA3 and AtCYP51G1. Positive control input showed that AtALA3 protein could be detected by myc antibody incubation, and AtCYP51G1 could be detected by flag antibody incubation, indicating the successful expression specificity of myc-AtALA3 and AtCYP51G1 flag in *N. benthamiana* leaves. CO-IP assays revealed that the AtCYP51G1-AtALA3 complex was specifically co-precipitated by anti-flag conjugated beads and detected by anti-myc immunoblotting only when both proteins were co-expressed, the complex was co-precipitated by anti-myc conjugated beads and detected by anti-flag immunoblotting only when both proteins were co-expressed (Fig. 5D), confirming their specific interaction.

While PCZ has been reported as a BR-specific inhibitor (*12*), and CYP51G1 catalyzes the production of campesterol-a direct precursor for BR biosynthesis (*29*), we investigated the functional link between the membrane target ALA3 and BR signaling. Phenotypic analysis of eBL-supplemented plants revealed that the *atala3* mutant exhibits a compact, dwarf architecture in adults and shortened roots in seedlings, a phenotype similar to that observed in both WT+PCZ and *atala3*+PCZ plants (fig. S8). While eBL application alone promoted growth and PCZ application alone suppressed it, the combined PCZ+eBL treatment relieved the inhibitory effect of PCZ (fig. S8A, C to E). This phenotypic difference was more pronounced in seedlings (fig. S8, B and F). This finding is consistent with the reported reference (*12*), which concluded that PCZ is a specific inhibitor of BR. Notably, eBL also restored the growth defects associated with *ALA3* deficiency (fig. S8). These findings indicate that AtALA3 engages in direct interaction with AtCYP51G1 and ALA3 may be involved in brassinosteroid biosynthetic pathway.

### Virtual screening-derived lead compounds demonstrated high-affinity binding to ALA3

To achieve pesticide reduction and efficacy enhancement beyond molecular design breeding, it is also possible to replace PCZ by designing compounds that target receptor proteins.We developed a virtual screening strategy targeting ALA3 active pocket using combined molecular docking (Vina) and 2D fingerprint similarity to PCZ. Screening 710,000 compounds from the ChemBridge database yielded 1,449 candidates, refined to 197 leads via ligand efficiency filtering, and ultimately 38 high-priority compounds based on structural features and pharmacophores (Fig. 6A). Docking energies between these compounds and ALA3 ranged from −10.87 to −7.96 kcal/mol, with classified structures: A01-A14/A17/A20/A21/A23/A25/A28/A30/A32/A34/A36 (amides), A15/A18/A33 (sulfonamides), and others (heterocycles) (fig. S9). Top-scoring representatives from each class (A01-amide, A15-sulfonamide, A22-heterocycle) were chemically synthesized (Fig. 6B and fig. S10 to 12).

**Fig. 6.**
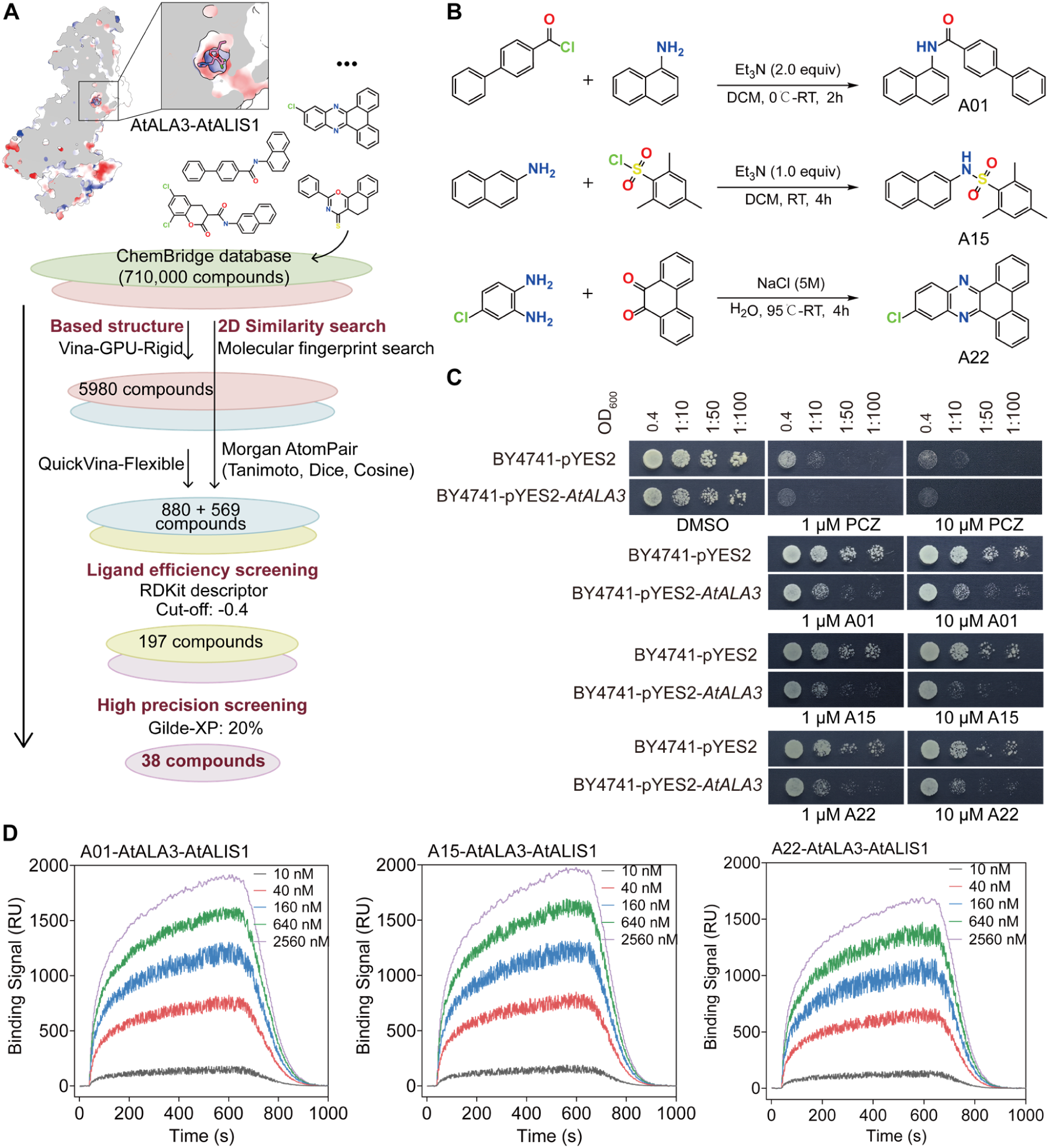
Computer-aided virtual screening and binding affinity assessment of potential ligands targeting AtALA3. (**A**) Schematic of virtual screening. Protein active pocket and 2D molecular fingerprints similarity-based virtual-screening strategy was performed, a database containing more than 710,000 compounds from ChemBridge was screened by Vina. PCZ was selected as representative search query molecules. 1449 candidates were selected for ligand efficiency screening, which depended on their structural features and physicochemical properties, with 38 hits found. (**B**) Synthetic routes for A01, A15, and A22 compounds. (**C**) Growth of BY4741 yeast strains supplemented homologous gene AtALA3 on SD/-His-Leu-Met medium containing DMSO, PCZ, A01, A15, or A22. Plates were photographed after 3 days of incubation at 30°C. (**D**) SPR affinity binding signals of AtALA3-AtALIS1 and A01/A15/A22 compounds.

Yeast-sensitivity assays of A01, A15, and A22 demonstrated that yeast strains expressing *AtALA3* showed significantly inhibited growth compared to pYES2-empty vector controls, with overexpression lines exhibiting more pronounced suppression than mutant complements. Notably, A01/A15/A22 treatments conferred greater tolerance than PCZ at equivalent concentrations (Fig. 6C and fig. S13). These results establish ALA3-sensitivity responsiveness to lead compounds A01, A15, and A22.

SPR assays quantified the affinity of A01, A15, and A22 for AtALA3, showing that SPR binding signals exhibited concentration-dependent enhancement for all three lead compounds (10 nM to 2560 nM) with AtALA3, generating characteristic interaction kinetic curves (Fig. 6D). The Avg KD for A01, A15, and A22 with AtALA3 were 5.11 × 10^-9^ M, 1.34 × 10^-8^ M, and 1.18 × 10^-7^ M, respectively, indicating strong binding affinities (table S4). Collectively, interactions of A01, A15, A22 and AtALA3 exhibits high-affinity binding, this rank order (A01>A15>A22) matched the virtual screening docking scores (fig. S9), validating the computational predictions.

In summary, virtual screening and experimental validation identified lead compounds A01, A15, and A22-designed to target the PCZ-binding pocket of ALA3, which suppressed growth in yeast heterologously expressing ALA3, exhibited high binding affinity to ALA3 protein.

### The lead compounds show growth-regulating effects and possess lower reproductive toxicity relative to PCZ

PCZ is widely used in agriculture, which may enter aquatic ecosystems and affect organisms. Early studies have demonstrated its reproductive toxicity in zebrafish (*Danio rerio*) models. The reported 96-h acute median lethal concentration (LC_50_) of PCZ to adult zebrafish is approximately 2 mg/L (*30–32*). Accordingly, we performed acute toxicity tests on adult zebrafish with the lead compounds. The results indicated that after 96 h of exposure, the mortality rate in the PCZ-treated zebrafish reached 36.67 ± 12.47%. Surviving individuals displayed clear signs of intoxication, including darkening of body color, food refusal, slowed and disoriented swimming, as well as lateral swimming and rolling. In contrast, neither mortality nor intoxication symptoms were observed in the solvent control or A01-treated groups, while the A15-treated group showed a low mortality rate of 3.33 ± 4.71%. Notably, all zebrafish in the A22-treated group died within 24 h of exposure (Fig. 7, A and B). The measurements of body length and weight revealed that, compared to the control group, zebrafish treated with PCZ or A22 showed significantly reduced body length and weight, along with delayed development. In contrast, no significant changes in these parameters were observed in the A01- or A15-treated groups, and their development proceeded normally (Fig. 7, C and D). Based on these findings, we conclude that, compared to PCZ, the lead compound A22 exhibits higher toxicity to aquatic organisms, whereas A01 and A15 are relatively safer.

**Fig. 7.**
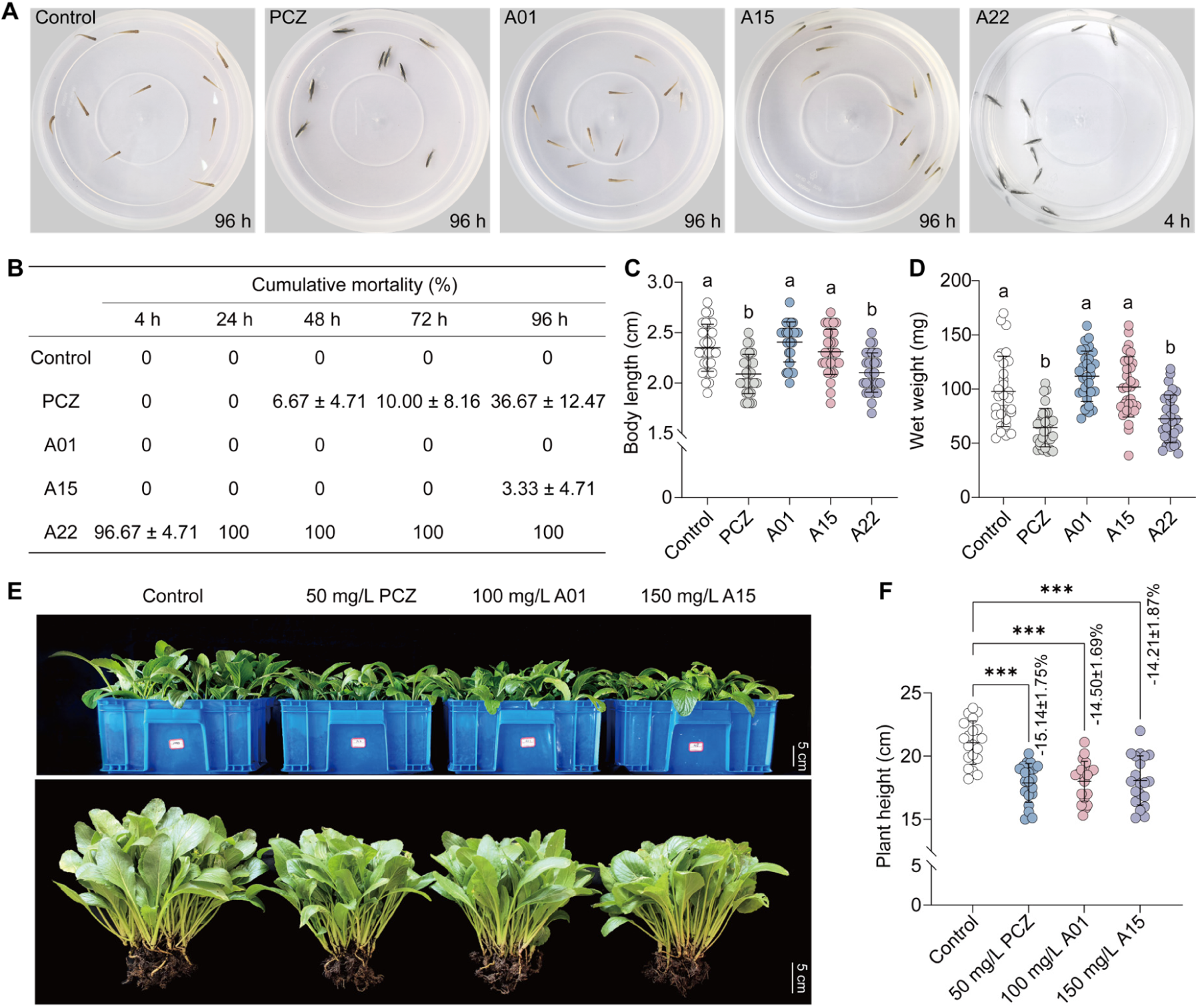
Assessment of reproductive toxicity and growth-regulating activity of the lead compounds. (**A**) Behavior of adult zebrafish (*Danio rerio*) in a toxicity assay with PCZ or lead compounds. (**B**) Cumulative mortality of adult zebrafish exposed to PCZ or lead compounds at 4, 24, 48, 72, and 96 h. Data are presented as the mean ± SD (*n* = 3, randomly assigned zebrafish with ten fish per treatment group). (**C and D**) Body length (C) and wet weight (D) of zebrafish at death across treatment groups. For zebrafish that succumbed to toxicity, measurements were taken at death; survivors were euthanized at 96 h and measured immediately. (**E**) 5 d phenotypes of DMSO, 50 mg/L PCZ, 100 mg/L A01, and 150 mg/L A15-treated *B. rapa*. Scale bar, 5 cm. (**F**) Comparison of plant height with PCZ/A01/A15-treated *B. rapa*. In (C) and (D), the error bars indicate the SD (*n* = 30). Letters indicate significant differences between mean values (one-way ANOVA: Tukey, *P* < 0.05). In (E) and (F), the error bars indicate the SD (*n* = 20). Asterisks indicate significant differences between mean values (one-way ANOVA: Dunnett, \*\*\**P* < 0.001). Percentages represent the inhibition rate of plant height.

To simulate practical application, we experimentally validated the growth-regulating efficacy of the lead compounds on the leafy *Brassica* vegetable, *Brassica rapa* var. *parachinensis* (*B. rapa*). Phenotypic analysis of *B. rapa* seedlings grown on 1/2 MS medium containing PCZ, A01, A15, or A22 demonstrated that all three lead compounds inhibited shoot and root growth compared to DMSO controls, though with reduced efficacy relative to PCZ. Compound potency followed the order A01 > A15 > A22 (fig. S14), consistent with both SPR binding affinities (table S4) and virtual screening docking scores (fig. S9). Furthermore, based on the zebrafish acute toxicity assays, the practical potential of lead compound A22 was ruled out. Following the results from the *B. rapa* seedling growth-regulation experiments, we further validated the efficacy of lead compounds A01 and A15 using pot-grown *B. rapa*. Building on the field application concentration of PCZ (50 mg/L), we increased the concentrations of the lead compounds, applying A01 at 100 mg/L and A15 at 150 mg/L. After 5 days of spraying, plant height was measured. Compared to the control group (DMSO), all three treatment groups (PCZ, A01, and A15) showed a significant reduction in *B. rapa* plant height, with inhibition rates of 15.14%, 14.50%, and 14.21%, respectively (Fig. 7, E and F).

In summary, this supports our research objective of designing receptor-targeting compounds to replace PCZ for growth regulation. The validation conducted to this point indicates that the lead compounds A01 and A15 possess promising potential for further optimization as next-generation pesticide candidates.

### Knockdown of *BraALA3a/b* reduced the sensitivity of *B. rapa* to PCZ

To better align with practical application, we conducted functional validation of *ALA3* homologs in *B. rapa*. Phylogenetic analysis identified three *ALA3* homologous genes in this species: *BraALA3a* (LOC103829275), *BraALA3b* (LOC103862629), and *BraALA3c* (LOC103843300). Among these, *BraALA3a* and *BraALA3b* share higher sequence similarity (fig. S3). Based on the *Arabidopsis* miR164a scaffold (*33*), we constructed an artificial microRNA (amiRNA) gene-silencing vector, pBI121-amiRNA-BraALA3a/b (Fig. 8A), designed to simultaneously knock down both *BraALA3a* and *BraALA3b*. The corresponding knockdown lines were subsequently generated in *B. rapa* via floral vacuum infiltration (*34, 35*). The genotyping results of the progeny showed that the expression levels of *BraALA3a* and *BraALA3b* were significantly reduced (Fig. 8, B and C), while the expression of *BraALA3c* showed a slight but non-significant decrease, confirming the successful generation of *BraALA3a/b* knockdown lines in *B. rapa*. Phenotypic analysis revealed that knockdown of *BraALA3a/b* markedly inhibited plant growth (Fig. 8D), leading to significantly reduced plant height, increased main stalk diameter, and decreased branch angles (Fig. 8, E to G). In addition, petiole length and leaf length were significantly shortened in the knockdown plants, whereas leaf width remained unchanged (Fig. 8, H to K). Compared with the wild type, the PCZ-induced inhibition rate of plant height was lower in the *BraALA3a/b* knockdown plants (21.66% vs. 45.53% in WT; Fig. 8E). While PCZ treatment increased the main stalk diameter by 25.29% in the wild type, no significant change was observed in the knockdown plants (Fig. 8F). Similarly, the inhibition rate of branch angle by PCZ was much lower in the knockdown plants (1.85%) compared to the wild type (22.60%; Fig. 8G). The knockdown plants also showed smaller inhibition rates in petiole length, leaf length, and total length of leaf upon PCZ treatment compared to the wild type (Fig. 8, H to J). These results indicate that knockdown of *BraALA3a/b* reduces the sensitivity of *B. rapa* to PCZ. This finding aligns with the observed resistance of the yeast *drs2* knockout strain to PCZ (Fig. 1) and the decreased sensitivity of the *Arabidopsis ala3* mutant to PCZ (Fig. 4). Collectively, the data demonstrate that *ALA3* homologs in *B. rapa* possess functional roles similar to those of *AtALA3* in *Arabidopsis*.

**Fig. 8.**
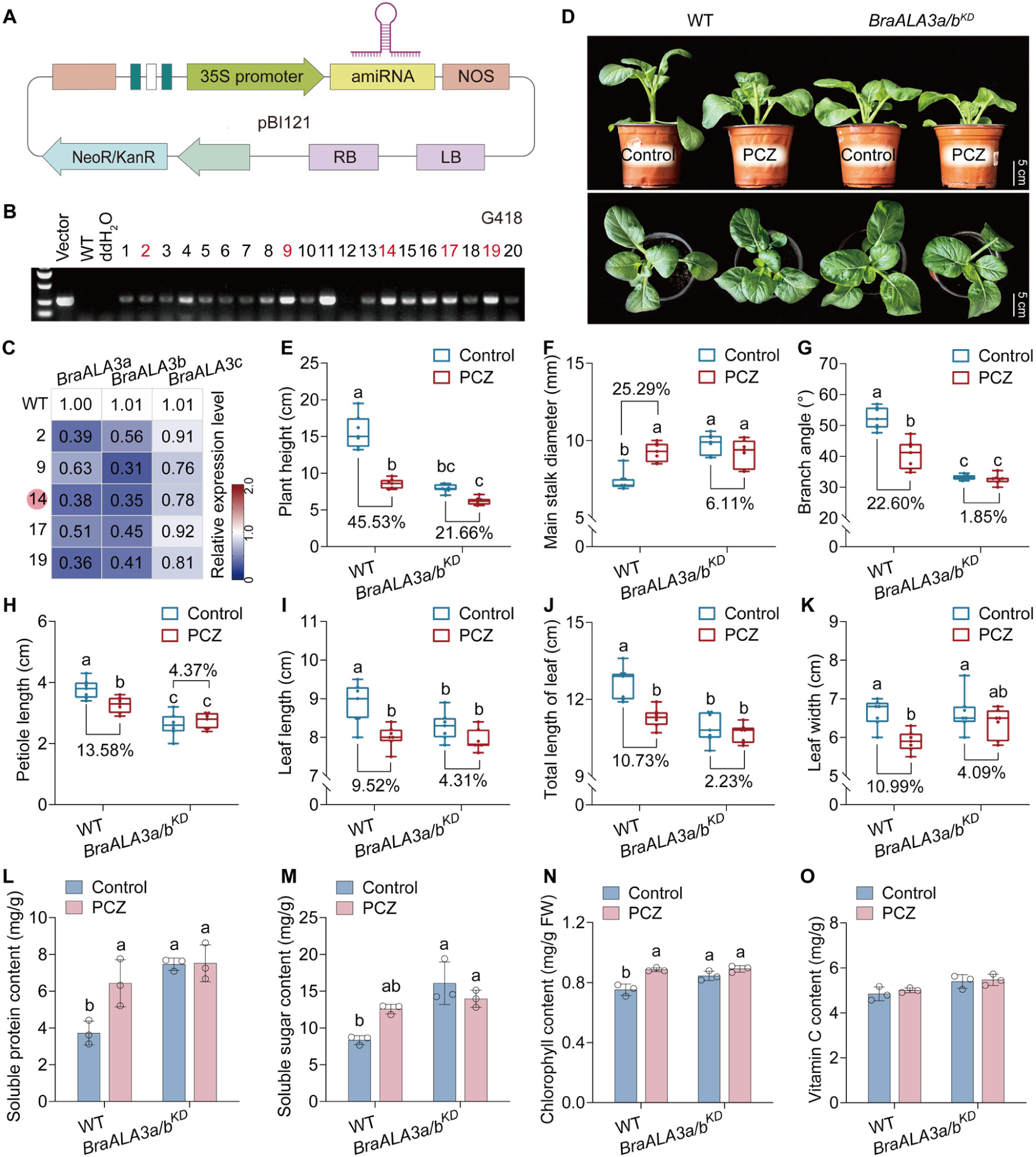
Generation and phenotypic analysis of *BraALA3a/b* knockdown lines in *B. rapa*. (**A**) Vector of floral dip transformation for generating *BraALA3a/b* knockdown lines (*BraALA3a/b^KD^*) in *B. rapa*. (**B**) Identification of *BraALA3a/b^KD^* by PCR. Plants labeled in red were selected for further analysis of gene expression. (**C**) Identification of *BraALA3a/b^KD^* by qRT-PCR. Expression profiles normalized to WT. Plants indicated by red circles were selected for phenotypic analysis. (**D**) 5 d phenotypes of control-treated or 50 mg/L PCZ-treated WT and *BraALA3a/b^KD^*. Scale bar, 5 cm. (**E** to **K**) Comparison of 5 d plant height (E), main stalk diameter (F), branch angle (G), petiole length (H), leaf length (I), total length of leaf (J), and leaf width (K) of control-treated or 50 mg/L PCZ-treated WT and *BraALA3a/b^KD^*. (**L** to **O**) Comparison of 5 d soluble protein content (L), soluble sugar content (M), chlorophyll content (N), and reduced vitamin C content (O) of control-treated or 50 mg/L PCZ-treated WT and *BraALA3a/b^KD^*. In (C), the data indicate the mean (*n* = 3). In (E) to (K), the whiskers indicate min to max (*n* = 7). In (L) to (O), the error bars indicate the SD (*n* = 3). Letters indicate significant differences between mean values (one-way ANOVA: Tukey, *P* < 0.05). Percentages represent the decrease or increase of each indicator.

Furthermore, physiological analysis of the *B. rapa* showed that the knockdown of *BraALA3a/b* led to elevated levels of soluble sugars, soluble proteins, and chlorophyll content (Fig. 8, L to N), while the content of reduced vitamin C showed no significant change (Fig. 8O). These results indicate that downregulating *BraALA3a/b* expression can enhance certain nutritional components in *B. rapa*. This finding holds significant implications for targeting ALA3 in leafy *Brassica* vegetables: editing ALA3 could enable the development of promising, PCZ-free varieties with improved growth regulation, not only modulating plant architecture but also potentially enhancing nutritional quality.

## Discussion

In field practices, PCZ is empirically applied by farmers to regulate plant architecture in leafy *Brassica* vegetables (*4–6*). This practice leads to PCZ accumulation and residue in plants, posing a threat to dietary safety (*1, 4, 5, 7*). Elucidating the molecular mechanism by which PCZ modulates plant growth is therefore of paramount importance. Although the sterol-to-brassinosteroid biosynthesis pathway has been identified as PCZ’s primary target (*10*), key gaps persist regarding how exogenously applied PCZ induces membrane remodeling and the mechanistic basis of its cellular uptake and trafficking to subcellular targets. In this study, we demonstrate for the first time that PCZ regulates plant growth by targeting the ALA3 membrane protein. Both *in vitro* and *in vivo* assays confirm their interaction. Moreover, we have identified compounds with the potential to serve as alternatives to PCZ. These findings open an important avenue for achieving green growth regulation in field applications.

Previous studies have established that ALA3 is a member of the phospholipid flippase P4-ATPase subfamily (ALAs). In the *Arabidopsis* genome, a total of twelve P4-ATPases have been identified, designated ALA1 through ALA12 (*36*). Our data indicate that, *atala3* mutants are still sensitive to PCZ even though the magnitude of difference between PCZ treated and untreated was smaller than that of WT (Fig. 4 and fig. S8). A similar phenotype was observed in knockdown lines of *BraALA3a/b* in *B. rapa* (Fig. 8). A plausible interpretation of these data is indeed that ALA3 is a target, but not the only one. Whether other members of the ALAs family might also interact with PCZ remains to be investigated. However, it has been reported that, with the exception of ALA2 (classified as P4C), all belong to the P4A-ATPase clade (*37*). Despite their close relationship, their substrate specificity, functions in regulating plant growth, and responses to abiotic stress are distinct. For instance, they differ in substrate preference: PS and phosphatidylethanolamine (PE) are the preferred substrates for *ALA3*, whereas *ALA5* and *ALA10* show a stronger preference for phosphatidylcholine (PC) (*21, 27, 38, 39*). They also differ in their growth-regulatory functions: *ALA3* specifically drives secretory vesicle generation in peripheral columella cells of the root tip (*21, 22*), *ALA4* and *ALA5* are primarily expressed in vegetative tissues, and their loss leads to defects in cell expansion (*39*). Their functions in responding to abiotic stress also differ: *ALA3* influences pollen tube tolerance to both high- and low-temperature stress (*40*), while *ALA1* and *ALA7* mediate the detoxification of fungal toxins in plants (*41*).

In summary, given the functional diversification within the ALAs family, the question of whether PCZ binds to other ALA proteins warrants future investigation. Such studies may help explain the residual sensitivity to PCZ observed in the *ala3* mutant. Importantly, our work provides strong evidence for a strong interaction between ALA3 and PCZ (Fig. 3, A to C). Supported by molecular docking, dynamics simulations (Fig. 2), and the resolved structure of the homologous dimer ScDRS2-Cdc50p (*20*), we propose a putative model in which PCZ inhibits ALA3 by competing with the lipid flippase regulator PI4P for the binding pocket within the transmembrane domain, thereby blocking the activation of its auto-regulatory conformational mechanism. This proposed model can subsequently be applied to validate drug transport using membrane protein-reconstituted liposomes or to screen for phospholipid flippase inhibitors. In previous studies, a range of emerging analogs, agonists, and inhibitors targeting phytohormone biosynthesis/signaling, and endomembrane trafficking, and so on have been well documented in both basic research and agricultural practice. For instance, pifithrin-α mimics ethylene by deactivating the ethylene receptor 1 (ETR1), thereby suppressing shade-induced hypocotyl elongation and associated gene expression (*42*). Our *in vitro* ATPase assays confirmed that PCZ inhibits ALA3 enzymatic activity (Fig. 3, D and E), providing experimental support for the molecular modeling-based proposal that PCZ likely acts as a competitive inhibitor of ALA3 (Fig. 9).

**Fig. 9.**
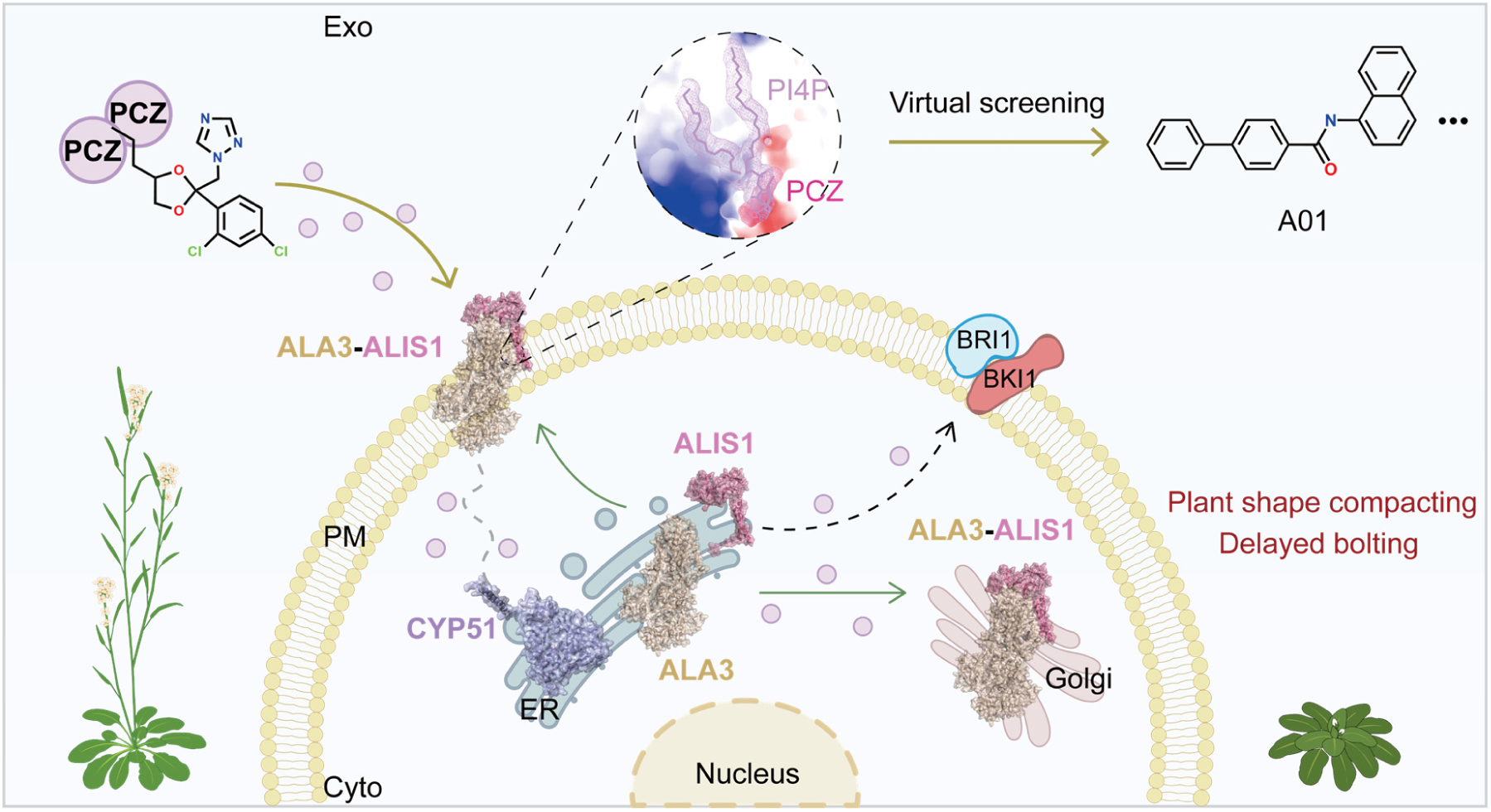
Mechanism model of PCZ targeting ALA3 to regulate plant growth and virtual screening of inhibitor leads. Green solid lines represent previously established mechanisms; gold solid lines indicate findings of this study; grey dashed lines denote incompletely resolved modes of action; black dashed lines correspond to mechanisms that remain uncharacterized. Based on the localization pattern that ALA3 localizes to the ER when expressed alone but relocates to the PM and Golgi upon co-expression with ALIS1 (*21, 22*), and given its interaction with CYP51, we propose a working model. Exogenously applied PCZ likely targets the membrane-localized ALA3-ALIS1 complex, potentially by competing with the lipid flippase regulator PI4P for its binding pocket. Following cellular entry, PCZ may then perturb BR biosynthesis and signaling through a series of intracellular events—potentially involving CYP51 and the Golgi-localized ALA3-ALIS1 complex, though the precise mechanism awaits full elucidation—thereby regulating plant growth. Lead compounds such as A01, which were screened by targeting this same binding pocket, exhibit potential as promising pesticide scaffolds. Their development could address the practical issue of PCZ misuse and abuse in the field.

ALA3 is not only a member of the extensive P4-ATP phospholipid flippase subfamily but also exhibits high evolutionary conservation across species (fig. S3 and Table S1). In this study, we confirmed the functional conservation of ALA3 homologs in *Arabidopsis* and *B. rapa*, which provides a potential editing target for exploring the roles of ALA3 orthologs in other species, such as monocots. Phylogenetic analysis revealed that ALA3 homologs from monocots cluster together and are phylogenetically more distant from DRS2, whereas those from dicots form clade located closer to DRS2 (fig. S3). This pattern suggests that functional divergence of ALA3 may have occurred between monocots and dicots, potentially leading to differences in specific biological roles. As a bolder speculation, precise editing of monocot *ALA3* genes could potentially generate semi-dwarf varieties of *Oryza sativa*, *Zea mays*, and *Triticum aestivum*, traits that are highly desirable in staple crops, while also addressing the issue of PCZ residue in the field (*12*). However, when pursuing such dwarfing traits, it is crucial to assess potential impacts on developmental fitness. Our study has shown that the *Arabidopsis ala3* mutant, although producing shorter siliques, remains fertile and capable of setting seeds (fig. S7) (*23*). This supports the feasibility of using ALA3 as a target for gene editing to develop dwarf, sturdy crop varieties. While the role of ALA3 in regulating pollen tube polarity during development has been documented (*23*), its potential involvement in other processes such as meiosis and gametophyte development remains unexplored. It is unknown whether PCZ controls the initiation of developmental programs like meiosis in a manner analogous to cyclin-dependent kinase (CDK) inhibitors (*43*). Moreover, whether editing the PCZ-interaction sites could be leveraged to select for improved crop varieties without compromising normal development awaits further investigation.

The specific intracellular processes affected by PCZ-mediated growth regulation through targeting the ALA3 membrane protein remain unclear. In fungi, the reported target of triazole pesticides is CYP51. Its plant homolog, the plant sterol 14α-demethylase CYP51G1 is an essential cytochrome P450 monooxygenase localized to the ER (*44*), which catalyzes the 14α-demethylation of sterol precursors (e.g., 24-methylene dihydrolanosterol) in the phytosterol biosynthesis pathway. Its catalytic product, campesterol, serves as the direct precursor for BR synthesis (*10, 44*). Thus, CYP51G1 activity indirectly modulates BR levels, thereby influencing plant growth, development, and environmental responses. Loss of *CYP51G1* function leads to accumulation of aberrant sterols and depletion of normal sterols, compromising membrane integrity, fluidity, and permeability, and ultimately impairing membrane protein function. These reports establish CYP51G1 within the genetic hierarchy of the BR pathway, upstream of the BR receptor BRI1 and its co-receptor BAK1 (*14, 45, 46*). Our study demonstrates a physical interaction between ALA3 and CYP51G1 (Fig. 6), suggesting these proteins may function coordinately in response to PCZ. However, the specific domains and key residues mediating their interaction remain undefined, as does the potential effect of PCZ on this protein complex (Fig. 9). And our research showed that *atala3* mutant phenotype defects were restored by eBL supplementation (fig. S8), coupled with prior evidence that BL reverses PCZ-induced growth suppression (*12*). These results collectively indicate that ALA3, when inhibited by PCZ, may influences BR-mediated pathway (Fig. 9). Within the membrane trafficking system, ALA3 acts as a central regulator whose ATP-dependent lipid flippase activity maintains the stability of the TGN-PM trafficking pathway (*21, 22*). The PM and ER dynamically exchange membrane components via vesicular transport, while the Golgi apparatus receives ER-synthesized proteins for further processing and sorting. A key question thus worth discussing is: what is the mode of the interaction between ALA3 and CYP51G1? One plausible scenario involves dynamic localization changes, while ALA3 primarily resides in the ER when expressed alone, its association with ALIS1 facilitates translocation to the PM and TGN (*21, 22, 27*). This redistribution may not be permanent or complete. Instead, the ALA3-ALIS1 complex might undergo partial or full dissociation after completing a round of vesicular transport, followed by reassembly upon initiation of the next transport cycle. During the temporal window between these cycles, ALA3, ALIS1, and CYP51G1 could coexist within the ER compartment, allowing their interaction. An alternative possibility is that, due to specific protein structural features, the ALA3-ALIS1 complex may stably localize to the PM and TGN once assembled. In this scenario, it could still interact with the ER-localized CYP51G1 via certain defined protein structural domains. This speculation is supported by documented cases in which two proteins residing in distinct subcellular compartments are still able to interact. For example, research has identified an ER-localized cyto-chrome *b_5_* (OsCYB5-2) that interacted with a high-affinity K^+^ transporter (OsHAK21) at PM (*47*). These insights guide future structural investigations into the ALA3-CYP51G1 interaction. Whether exogenous PCZ application alters the ALA3-ALIS1 interaction and subcellular localization, whether ALA3 subsequently reverts to ER localization, and how such dynamic redistribution might affect the association between ALA3 and CYP51G1 (Fig. 9) remain open questions. These proposed mechanisms represent plausible but currently unvalidated models that will require further investigation using approaches such as real-time live-cell imaging and rigorous membrane fractionation assays.

The rapid advancement of virtual screening technologies in comparative genomics has bridged computational chemistry and molecular biology, emerging as a powerful interdisciplinary tool. Recent applications in pesticide development include: *Striga* germination is induced by host-root-derived strigolactone (*48*), the identification of *(S)-4a*, a strigolactone agonist targeting ShHTL7 through virtual screening, which acts as a suicidal germination inducer to control root parasitic weeds (*49*); and the discovery of small-molecule inhibitors (ligand_SL2 and ligand_3275) that disrupt Wheat dwarf virus (WDV) Rep protein activity, effectively suppressing viral replication (*50*). After confirming the interaction between PCZ and ALA3, we targeted the PCZ-binding pocket of ALA3 to screen for potential inhibitors, and verified that the lead compounds exhibit high binding affinity to the ALA3 protein (Fig. 6, C and D; fig. S13; table S4). While these compounds do possess growth-regulating activity, their efficacy is lower than that of PCZ (Fig. 7, E and F; fig. S14). This difference can be attributed to several factors. The lead compounds identified by virtual screening are initial, single-target hits with modest efficacy, unlike commercial pesticides which are optimized for multi-target synergy. Their unoptimized pharmacokinetic and physicochemical properties contrast with the extensively refined scaffolds of commercial pesticides. Additionally, leads are tested as pure compounds, whereas commercial pesticides include adjuvants that enhance permeability, stability, and on-target deposition. Furthermore, the screening classified candidates into amides, sulfonamides, and heterocycles, with representative leads synthesized and validated. The many remaining candidates also hold potential value. Therefore, given these inherent limitations of the lead compounds and inspired by relevant references (*49, 51*), future work will focus on systematic structural optimization. This will involve synthesizing a series of analogs based on the current scaffold to delineate the structural moieties critical for activity, elucidate key pharmacophores, improve pharmacokinetic properties, and develop formulation. Through these approaches, the initial leads can be methodically refined and upgraded.

In summary, our study demonstrates that PCZ modulates plant growth through targeted inhibition of the membrane flippase ALA3. We elucidate PCZ’s mechanism of action in disrupting phospholipid translocation, redefine its growth-regulatory paradigm beyond conventional single-target pesticide models, and propose a cross-regulatory network linking PCZ’s effects on ALA3, its subunit ALIS1, and sterol 14α-demethylase (CYP51G1). These findings provide both a molecular target for breeding PCZ-free cultivars via genome editing, and lead scaffolds for developing ALA3-targeted pesticide with improved selectivity. This work bridges plant chemical biology and rational pesticide design, offering transformative strategies for sustainable agriculture.

## Materials and Methods

### Chemicals and materials

The PCZ active compound and the starting materials for lead compound synthesis were purchased from Shanghai Bepharm Science & Technology Co., Ltd. The eBL (24-eBL) (*52*) were purchased from Shanghai yuanye Bio-Technology Co., Ltd. The lead compounds, designated A01, A15, and A22 following structural validation by nuclear magnetic resonance (NMR) spectroscopy, were advanced for functional characterization. Other reagents were purchased from Sigma-Aldrich Chemical Corporation, Ltd. (St. Louis, MO, USA). Primers were synthesized by Tsingke Biotechnology Co., Ltd. (Beijing, China) (table S2). Yeast library was generously provided by Takashi Akiihiro, Shimane University, Matsue, Japan. The *Saccharomyces cerevisiae* BJ5465 strain was obtained from American Type Culture Collection (ATCC). The pRS424 and pRS426 vectors were generously provided by Long Li, Peking University, Beijing, China. Other vectors were kept in our laboratory.

### Plant materials and growth conditions

All *A. thaliana* materials were in the Columbia-0 (Col-0) ecotype background. T-DNA insertion lines *atala3-p* (SALK_129494C) and *atala3-l* (SALK_205861C) were purchased from Arashare (https://www.arashare.cn/), while *atala3-4* (SALK_082157) homozygous seeds were generously provided by Yun Xiang (*23*), with hygromycin (Hyg) resistance and genotyping confirming their identity. For pot experiments, *A. thaliana* seeds were stratified at 4°C for 2 days before sowing on sterilized substrate, with plants grown at 20-22°C, 70% humidity, 150 µmol·m^-^²·s^-^¹ light intensity (16/8 h photoperiod), and watered with 0.1% nutrient solution every 3-5 days. For aseptic seedling growth assay, surface-sterilized *A. thaliana* seeds were plated on 1/2 MS medium (1/2 MS, 3% sucrose, pH 5.8, 4 g/L agar), 4°C, darkness for 2 days, then grown at 22°C for 4 days before chemical treatment.

*B. rapa* materials were of the ‘Youqing Sijiu’ ecotype background. For aseptic seedling growth assay, sterilized seeds were plated on 1/2 MS medium, 25°C, dark-germinated for 12 h, then grown for 2 days under controlled conditions (25°C, 70% humidity, 150 µmol·m^-^²·s^-^¹, 16/8 h photoperiod) prior to chemical treatment.

The study utilized *N. benthamiana* plants, with seeds sown directly into sterilized substrate under conditions identical to those used for *A. thaliana* pot cultivation.

### PCZ/eBL/lead compound treatment and phenotype analysis

Potted *A. thaliana* plants were treated with PCZ at 28 days post-sowing, with petiole length, leaf length, and leaf width measured in mature leaves post-treatment. The rosette leaf arrangement in *A. thaliana* refers to previously reported literature (*53*). For aseptic seedling growth assay, 4-day-old *A. thaliana* seedlings were transferred to 1/2 MS medium containing PCZ, and root elongation was quantified after 5 days of exposure. For the combined PCZ and eBL treatment, the compounds were pre-mixed prior to application.

Potted *B. rapa* plants were treated with PCZ or the lead compounds prior to bolting, and the post-treatment phenotypes were recorded 5 days after application. For aseptic seedling growth assay, 2-day-old *B. rapa* seedlings were transferred to 1/2 MS medium containing PCZ, with hypocotyl and root elongation measured after 4 days.

### Screening of PCZ sensitive genes by yeast library

The gene-knockout strain was cultivated into a 96 deep-well plate containing 200 μL YPD liquid culture medium (2% polypeptone, 1% yeast extract, 0.04% adenine, 2% glucose, pH 5.8) at 30°C for 12 h. The OD_600_ of cultures were measured by a microplate reader and diluted to 0.01 by sterile water. The 3 μL diluted yeast fluid was spotted on YPDA solid culture medium (2% polypeptone, 1% yeast extract, 0.04% and adenine, 2% glucose, 2% agar, pH 5.8) containing including 0.5 μM, 1 μM, or 10 μM PCZ. After 2-3 d, the plates were observed and YPDA solid culture medium containing DMSO as a control (*25, 26*).

### Analysis of phylogenetic tree

The DRS2 (YAL026C) sequence was searched on NCBI (https://www.ncbi.nlm.nih.gov/) and blasted for homologous genes, multiple protein sequences were downloaded and phylogenetic tree analysis was performed using Poisson model based on neighbor-joining (NJ) method in MEGA11.0 software to create Clusterw multiple sequence alignments (*54*).

### Heterologous expression in yeast

Total RNA of *A. thaliana* was extracted with the E.Z.N.A. Plant RNA Kit (Omega, Guangzhou, China), and reverse transcription was conducted with the PrimeScript^TM^ RT reagent Kit and gDNA Eraser (Takara, Dalian, China). The CDS region of *AtALA3* was amplified by 2 × Primer STAR Max premix (Takara), *AtALA3* was subcloned into pYES2 by ClonExpression^Ⓡ^Ⅱ One Step Cloning Kit (Vazyme, Nanjing, China). Primers were shown in Supplementary Tab. 2. The pYES2-*AtALA3* were supplemented in *drs2* yeast or BY4741 (*MATa his3-1 leu2-0 met15-0 ura3-0*). The preparation and transformation of yeast receptive state were carried out according to Yeastmaker™ Yeast Transformation System 2 (Takara), and positive transformants were identified using Quick Yeast positive clone assay Kit (Coolaber, Beijing, China).

Complement transformants and enriched expression transformants were incubated in 500 μL SD/-His-Leu-Met medium (0.67% YNB, 2% D-Galactose, 0.002% His, 0.002% Met, 0.01% Leu, pH5.7) for 12 h at 210 rpm, 30℃. Then, transformants were replaced with fresh culture medium and cultured for 4 h, OD_600_ was adjusted to 0.1, 0.01 with distilled water (ddH_2_O). 3 μL transformants diluted fluid were spotted onto SD/-His-Leu-Met medium (2% Agar) containing DMSO, 0.5 μM or 1.0 μM PCZ (or lead compounds), and cultured at 30°C for 2-3 d (*55*). Each experiment was repeated independently at least 3 times.

Complement transformants and enriched expression transformants were incubated in 3 mL SD/-His-Leu-Met medium overnight at 30℃. OD_600_ was adjusted to 0.04 with corresponding fresh culture medium and DMSO/0.5 μM PCZ. Yeast cells were induced to express at 210 rpm, 30℃, and OD_600_ was measured at 9, 12, 15, 18, 21, 24, 33, 36, 39, 42, 45, 48, 60 h and 72 h for growth curve plotting (*25, 26*). Each experiment was repeated independently at least 3 times.

### Protein 3D modelling of AtALA3-AtALIS

A homology model of the AtALA3-AtALIS1 complex was generated based on the structure of the DRS2-Cdc50p template (*20*). The amino acid structures were obtained from the UniProt database (https://www.uniprot.org/), and were hosted on the Google Colaboratory project hosting platform using ColabFold, the structure of the polypeptide chain complexes was predicted by multichain mode, with other parameters defaulted. After submitting the task, ColabFold was trained using transformer model and outputed a predicted structure file (PDB format) (*56*). The predicted model was reviewed by PyMOL visualisation software. The complex model was evaluated and optimised using the methods of Ramachandran (*57*). A structural model of the AtALA3-AtALIS1 complex was generated using this approach (fig. S4A). This structural feature is consistent with typical P-type ATPase family members (*58*). After the optimisation, no residues were found in the restricted region and the overall concentration of the main chain dihedral angle was towards the most acceptable region (fig. S4, B and C), indicating that the conformational rationality of the structure has been significantly improved, and this structure was used for molecular docking of subsequent protein models (Method S1).

### Molecular docking of AtALA3-AtALIS with PCZ

Based on previous reports (*20, 59*), the parameters of the docking box were determined (box dimensions: 30 Å × 30 Å × 30 Å, lattice length: 0.375 Å). Molecular docking was performed using Autodock Vina (*60*), vina.exe was used for docking and vina_split.exe was used to split the docking results, exhaustiveness = 100 was setted for more accurate results. The ligand-receptor interactions in each binding mode, including hydrogen bonding, hydrophobic forces, and van der Waals forces between the ligand and active site residues, were analysed by PyMOL and UCSF ChimeraX (Method S2).

### Molecular dynamics simulation (MD)

50 ns all-atom molecular dynamics simulations of the complex model were performed using the GROMACS software package (*61*). The molecular dynamics trajectories were used to analyse the structure, conformational dynamics and interaction characteristics of protein-small molecule complexes. The stability of conformations was assessed by RMSD, RMSF, and Rg (Method S3).

### Site-directed mutagenesis verification

The amino acid properties were changed according to the principle of polar-nonpolar and acidic-basic changes (Table. S3), the amino acid residues constituting the PCZ-ALA3 binding interface were targeted for mutation. Primers for site-directed mutagenesis were designed through the Quik Change Primer Design online website (https://www.agilent.com.cn/store/primerDesignProgram.jsp) and site-directed mutagenesis pYES2-AtALA3 vector was constructed using the Fast Site-Directed Mutagenesis Kit (TIANGEN, Beijing, China), which was supplemented with the gene knockout yeast *drs2*, and yeast-sensitive of PCZ was performed (*26*).

### P4-ATPase protein expression

Homologs of ALA3 and their *β*-subunits from yeast and *A. thaliana* were cloned into pRS426 (sfGFP-TwinStrep-3C-CtP4-ATPase, Ura resistance selection) and pRS424 (sfGFP-His-3C-β-subunit, Trp resistance selection) vectors, respectively, then co-transformed into BJ5465 (*MATa ura3-52 trp1 leu2-delta1 his3-delta200 pep4::HIS3 prb1-delta1.6R can1 GAL*). Transformants were selected on SD-Ura/Trp medium (6.7 g/L YNB with ammonium sulfate, 1.29 g/L DO supplement-Ura/Trp, 2% glucose, pH 5.7) and verified by colony PCR. Selected clones were cultured in SD-Ura/Trp with 2% raffinose (30°C, 220 rpm, 24 h) until OD^600^ reached 5, then induced with 4×YP medium (40 g/L yeast extract, 80 g/L peptone, 8% galactose) at 25°C, 220 rpm, 20 h. Cells were harvested by centrifugation (4°C, 3,000 × g, 20 min) for protein extraction.

Membrane proteins were extracted through differential centrifugation and detergent solubilization. Yeast cells were resuspended in 2 volumes of lysis buffer (20 mM Tris-HCl [pH 7.4], 150 mM NaCl, 5 mM MgCl_2_, 1 mM DTT, 1:500 protease inhibitor cocktail), disrupted by high-pressure homogenization (1,800 bar, 5-7 passes), and clarified by centrifugation (4°C, 20,000 × g, 25 min). Membrane fractions were collected via ultracentrifugation ((4°C, 44,000 rpm, 1 h, 2 passes), then solubilized in 5 volumes of extraction buffer (20 mM Tris-HCl [pH 7.4], 10% [v/v] glycerol, 150 mM NaCl, 5 mM MgCl_2_, 2% lauryl maltose neopentyl glycol [LMNG], 1 mM DTT, 1:500 protease inhibitor cocktail) for 1 h at 4°C. The solubilized fraction was recovered by ultracentrifugation (4°C, 44,000 rpm, 1 h) for subsequent analyses. Protein expression and purity were assessed by SDS-PAGE, with concentration determined using Enhanced BCA Protein Assay Kit (Beyotime, Shanghai, China) (*62*) (fig. S5).

### Binding affinity determination by SPR

Protein-ligand interactions between ScDRS2-ScCdc50/AtALA3-AtALIS dimers and compounds were analyzed by SPR. Compounds were immobilized on 3D photo-crosslinked chips using a Biodot™ AD1520 array spotter, followed by vacuum drying, UV crosslinking, and sequential washing (DMF, ethanol, H_2_O; 15 min each) before nitrogen drying and Flowcell Cover assembly. Detergent/glycerol-free protein samples (PBS-dialyzed and concentrated) were diluted, and diluted in PBST (pH 7.4, 0.1% Tween 20) at five concentrations (10, 40, 160, 640, 2560 nM). Samples were injected in ascending concentration order (0.5 μL/s, 4°C, 600 s association/360 s dissociation), with regeneration using glycine-HCl (pH 2.0, 2 μL/s) (*63*). Kinetic parameters (Ka, Kd) and equilibrium dissociation constants (KD) were derived from binding curves.

### *In vitro* ATPase activity assay

Functional validation of protein complex bioactivity was performed through *in vitro* ATPase activity assays, including testing for potential P4-ATPase activators or inhibitors. Using a Malachite Green Phosphate Assay Kit (Sigma), liberated phosphate from ATP hydrolysis was quantified colorimetrically at 620 nm with or without POPS, with standard calibration curves enabling precise activity quantification across experimental conditions. POPS was prepared as 10 mM stock in 1% C_12_E_9_ solution, while ATP (12.5 mM stock) was aliquoted and stored at −80°C. Protein complexes were pre-incubated with PCZ for 1 h prior to ATP addition, with BeF3^-^ (a known P-type ATPase inhibitor) serving as negative control (*62*). All experimental conditions were independently replicated three times.

### Histological section

The samples were embedded by Eponate 12™-Araldite embedding Kit with DMP-30 (TED PELLA, INC). The samples were polymerized at 60℃ for 16-24 h until the samples cooled to certain hardness. The samples were trimmed to the appropriate size and section, embedded plant tissues were sectioned by the *Leica RM2235* manual microtome in 5-10 μm. The toluidine blue-stained sections were observed under Laser capture micro-dissection (LMD) (*25*). Cell length and width were quantified from stained samples using Image-Pro Plus 6.0 software (Method S4).

### Quantitative real-time PCR (qRT-PCR)

Total RNA was extracted from PCZ-treated *A. thaliana* samples followed by reverse transcription. RT-qPCR was performed using ChamQ SYBR qPCR Master Mix (Vazyme, Nanjing, China). *Actin* served as the reference gene for *A. thaliana* (*64*). All experiments were performed with 3 independent biological replicates. The Ct value from the BioRad CFX96TM Real Time PCR System were derived, the expression level of the target genes was calculated under different treatments using equation 2^-ΔΔCt^ (the expression level of the internal reference gene is 1). The primers used for RT-qPCR analysis are listed in table S2.

### Yeast two-hybrid (Y2H) assay

Protein-protein interactions were validated via yeast two-hybrid (Y2H) assays. Candidate proteins were cloned into AD (activation domain) and BD (binding domain) vectors, transformed into NMY51 yeast (*MATa, his3Δ200, trp1-901, leu2-3,112, ade2*), and tested for autoactivation. The minimal 3-amino-1,2,4-triazole (3AT) concentration suppressing autoactivation was determined before co-transforming candidate proteins. Transformants were plated on selective media SD-TLH + X-gal (26.7 g/L Minimal SD Base, Do-Supplement-Trp/Leu/His, pH 5.8, 30 mM 3AT, 40 µg/mL X-gal) and SD-TLHA + X-gal (26.7 g/L Minimal SD Base, Do-Supplement-Trp/Leu/His/Ade, pH 5.8, 30 mM 3AT, 40 µg/mL X-gal), with growth recorded after 3 days at 30°C (*47*).

### Bimolecular fluorescence complementation (BiFC)

pCAMBIA1300-YFPN-AtALA3 and pCAMBIA1300-AtALIS1 was transiently co-expressed with pCAMBIA1300-AtCYP51G1-YFPC (or pCAMBIA1300-YFPN-AtALA3 + pCAMBIA1300-AtALIS1/pCAMBIA1300-AtCYP51G1-YFPC with the empty vector as the control) in *N. benthamiana* leaves. LSCM analysis (excitation wavelength of 514 nm) was performed 60 h after transfection (*22*).

### Luciferase complementation assay (LCA)

pCAMBIA1300-AtALA3-CLuc and pCAMBIA1300-ATALIS1 was transiently co-expressed with pCAMBIA1300-NLuc-AtCYP51G1 (or pCAMBIA1300-AtALA3-CLuc + pCAMBIA1300-ATALIS1/pCAMBIA1300-NLuc-AtCYP51G1 with the empty vector as the control) in *N. benthamiana* leaves. *N. benthamiana* leaves were collected at 60 h post infiltration. Luciferase activity was assayed by infiltrating D-luciferin potassium salt solution (YEASEN) into inoculated *N. benthamiana* leaves, followed by 10 min of dark incubation on ice. Luminescence was captured using Image Lab software (*65*).

### Co-immunoprecipitation (Co-IP)

AtALA3 (tagged with MYC at N-terminus), AtALIS1, and AtCYP51G1 (tagged with 3×FLAG at N-terminus) were cloned and constructed into the pCAMBIA1300 vector, and transformed into GV3101-p19. *N. benthamiana* leaves transiently co-expressing myc-AtALA3, AtALIS1, and AtCYP51G1-flag, or myc-AtALA3+AtALIS1/AtCYP51G1-flag with the empty vector (as the control), were collected at 60 h post infiltration. Total proteins were extracted from homogenized *N. benthamiana* leaves in lysis buffer (Beyotime) containing 1 mM PMSF. After solubilization by vortexing and incubation at 4°C for 30 min, the protein extracts were cleared at 14,000×g for 5-20 min. 4 µg MYC/FLAG monoclonal antibody (ABclonal, Wuhan, China) and 300 µL Incubation buffer (0.2 g/L KCl, 0.2 g/L KH_2_PO_4_, 1.14 g/L Na_2_HPO_4_ꞏ12H_2_O, 8 g/L NaCl, 0.42 g/L NaF, 1.86 g/L EDTA·2Na·2H_2_O, pH 7.4) was added to the 3 mg protein lysates at 4°C overnight with gentle shaking. Reaction mixture was incubated with 100 µLProtein A/G beads (Proteintech, Wuhan, China) by low-speed centrifugation in PBS at 4°C for 4 h with gentle shaking. Immunoprecipitated complexes were washed five times with 1×Washing buffer (2.42 g/L Tris, 8.76 g/L NaCl, 0.373 g/L KCl, 2 mL/L Tween-20, pH 7.4). Proteins bound on the beads were boiled at 75℃ for 10 min with 5 × SDS loading buffer and analyzed by immunoblotting. Actin was used as a loading control, with Input lysates serving as the experimental reference (*22, 47*).

### Virtual-screening

A ChemBridge database contained 710,000 commercial compounds that were used for the screening. Large-scale virtual screening targeting the ALA3-PCZ binding pocket was performed using Vina-GPU-Rigid for primary docking, followed by refinement with QuickVina-Flexible. Candidate compounds were subjected to 2D similarity searches based on Morgan and AtomPair fingerprints (Tanimoto, Dice, Cosine), then filtered by RDKit descriptors (cutoff: −0.4) to eliminate structurally complex or low-efficiency molecules. Top-ranked hits underwent structure- and pharmacophore-based prioritization using Glide XP (top 20%), yielding final lead compounds (*49*).

### Lead compound synthesis

Synthesis of A01: In a 30 mL two-necked round-bottom flask equipped with a magnetic stirrer under argon atmosphere, naphthalen-1-amine (5.0 mmol, 1.0 equiv.) and triethylamine (4.0 mmol, 2.0 equiv.) were dissolved in anhydrous dichloromethane (DCM, 4 mL). A solution of the corresponding 4-phenylbenzoyl chloride in anhydrous DCM (4 mL) was added dropwise to the mixture at 0°C. After warming to room temperature, the reaction was stirred for 2 h until complete consumption of the aniline (monitored by TLC). The mixture was quenched with 1N HCl (10 mL) and extracted with DCM (2 × 10 mL). The combined organic layers were washed with brine, dried over sodium sulfate, and concentrated under reduced pressure. The residue was recrystallized from boiling tetrahydrofuran to afford N-(naphthalen-1-yl)-[1,1’-biphenyl]-4-carboxamide (*66*).

Synthesis of A15: In a 20 mL round-bottom flask, *β*-Naphthylamine (1.0 mmol) and 2-mesitylenesulfonyl chloride (1.0 mmol) were dissolved in dichloromethane (10 mL), followed by dropwise addition of triethylamine (0.1 mmol). The reaction proceeded at room temperature for 6 h, after which the resulting benzenesulfonamide derivative was purified by column chromatography using ethyl acetate-petroleum ether (1:5 v/v) as the eluent (*67*).

Synthesis of A22: In a 20 mL round-bottom flask, 9,10-phenanthrenedione (1.0 mmol) and 4-chlorobenzene-1,2-diamine (1.1 mmol) were dissolved in ethanol (10 mL) and stirred under reflux at 70°C. After 30 min of reaction, a significant increase in suspended solid volume was observed, prompting continued stirring for an additional 2 h. The liquid quinoxaline derivative was subsequently purified by column chromatography using ethyl acetate-petroleum ether (1:2 v/v) as the eluent (*68*). All synthesized compounds were verified for purity by NMR.

### Zebrafish acute toxicity assay

Zebrafish acute toxicity assay was conducted with the lead compounds. Following the Test Guidelines on Environmental Safety Assessment for Chemical Pesticides (GB/T 31270.12-2025) issued by the Ministry of Agriculture of China, a semi-static method was employed. The solvent served as the positive control, 2 mg/L PCZ as the negative control (*31, 32*), and 2 mg/L A01, A15, and A22 as experimental groups, with three replicates per treatment. Signs of intoxication and mortality in zebrafish (2-month-old) were monitored and recorded at 4, 24, 48, 72, and 96 hours after exposure to the compounds (*30*).

### Knockdown of BraALA3a/b in B. rapa

Based on the backbone of *Arabidopsis* miR164a (*33*), an artificial microRNA (amiRNA) gene-silencing vector targeting both *BraALA3a* and *BraALA3b* was constructed and designated pBI121-amiRNA-BraALA3a/b (Fig. 8A). *B. rapa* plants at the early flowering stage were transformed via vacuum infiltration (*34, 35*), followed by 6-8 rounds of manual pollination using untransformed pollen (Method S5). Seeds were harvested after maturation. *B. rapa* plants (28-day-old) were treated with PCZ, and the phenotypes were recorded 5 days after application.

### Statistical analysis

Data were presented as means ± standard deviations (SD). Based on different comparisons, significant differences were calculated by two-way ANOVA (Dunnett or Šídák, \**P* < 0.05, \*\**P* < 0.01, \*\*\**P* < 0.001, no mark presents no significance), or one-way ANOVA (Tukey, *P* < 0.05). Statistical graphs were drawn using GraphPad Prism 10.3.

## Acknowledgments

We thank T. Akihiro (Faculty of Life and Environmental Science, Shimane University) for providing the yeast library, Y. Xiang (School of Life Sciences, Lanzhou University) for supplying mutant seeds, and L. Li (School of Life Sciences, Peking University) for providing protein expression vectors. We also thank B. Liu and X. Wu (College of Materials and energy, South China Agricultural University) for technical support in virtual screening, as well as G. Yao and H. Gao (College of Plant Protection, South China Agricultural University) for assistance in compound synthesis.

## Funding

Key-Area Research and Development Program of Guangdong Province (2022B0202080001)

Generic Technique Innovation Team Construction of Modern Agriculture of Guangdong Province (2023KJ130)

## Author contributions

Conceptualization: Q.G., F.L., and H.X.

Methodology: Q.G., S.W., D.G., Z.C., and X. Z.

Investigation: Q.G., Y. S., and Y.Y.

Visualization: Q.G., S.W., and R.C.

Supervision: F.L. and H.X.

Writing—original draft: Q.G. and R.C.

Writing—review & editing: Q.G., R.C., H.X., and F.L.

## Competing interests

Authors declare that they have no competing interests.

## Data, Code, and Materials Availability

All data and code needed to evaluate and reproduce the results in the paper are present in the paper and/or the Supplementary Materials. This study did not generate new materials.

## Supplementary Materials

This PDF file includes:

Figs. S1 to S14

Tables S1 to S4

Methods S1 to S5

## Notes

### Competing Interest Statement

The authors have declared no competing interest.

