## supplementary_materials for "Targeting ALA3 with propiconazole regulates plant growth and enables discovery of promising inhibitor leads"

Qing Gao *et al.*

**This PDF file includes:**

Supplementary Text
Figs. S1 to S14
Tables S1 to S4
Methods S1 to S5

### Supplementary Text

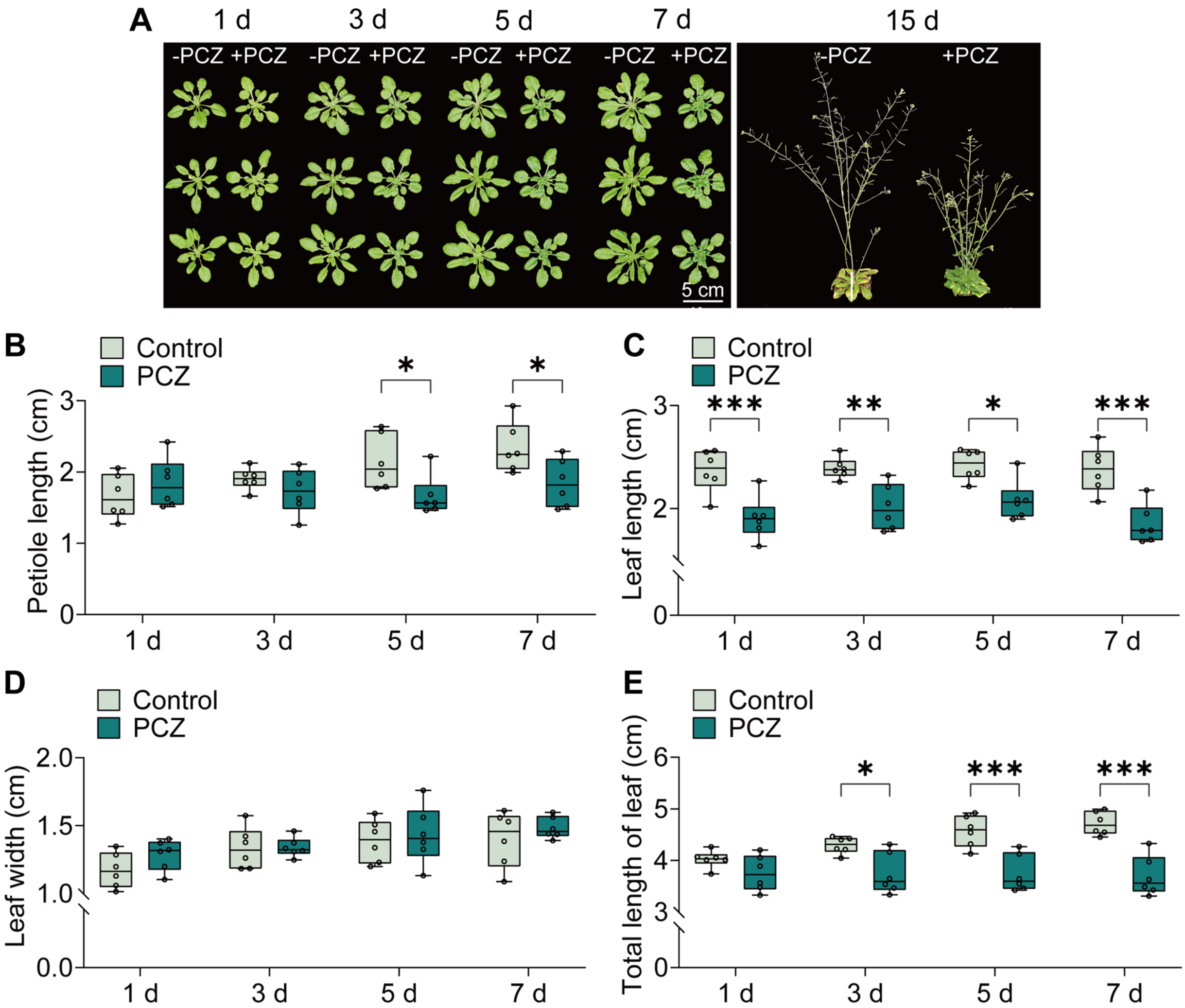


**Fig. S1.**

**Growth inhibition of *Arabidopsis thaliana* (*A. thaliana*) by PCZ. (A)** 1 d, 3 d, 5 d, 7 d, and 15 d phenotypes of control-treated or 50 mg/L PCZ-treated *A. thaliana*. Scale bar = 5 cm. **(B-E)** Comparison of 1 d, 3 d, 5 d, and 7 d petiole length (B), leaf length (C), leaf width (D), and total leaf length (E) of control-treated or 50 mg/L PCZ-treated *A. thaliana* mature leaves (L6-8). The rosette leaf arrangement in *A. thaliana* refers to previously reported literature, *Hunziker et al. (53)*. In **B-E**, the error bar indicates the SD ($n = 6$). Asterisks indicate a significant difference between mean values of control and PCZ treatment (two-way ANOVA: Šídák, $*P < 0.05$, $**P < 0.01$, $***P < 0.001$, no mark indicates no significance).

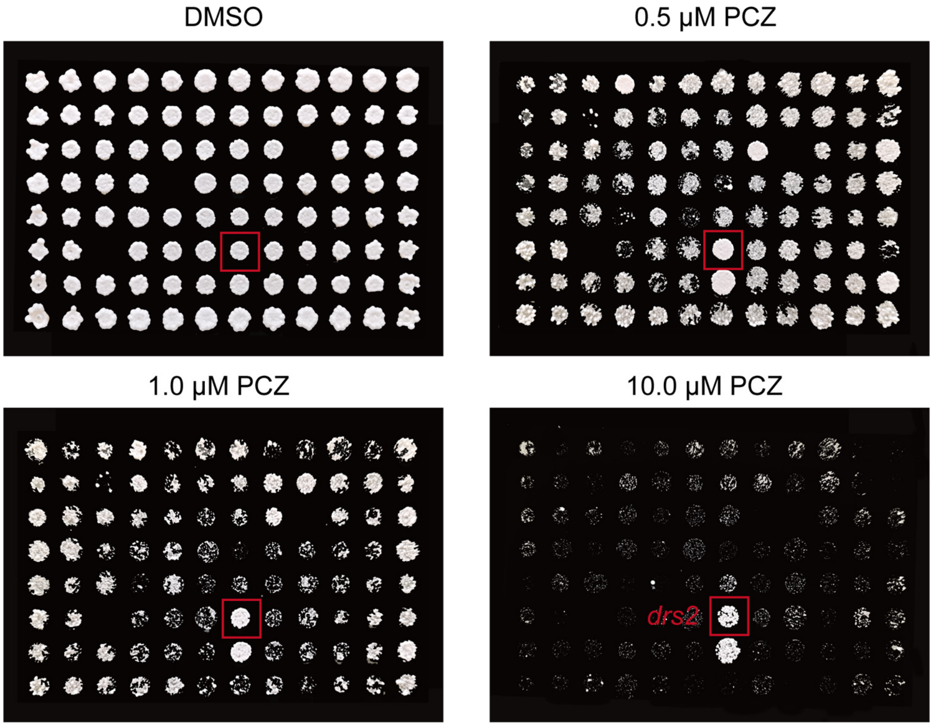


**Fig. S2.**

**Identification of PCZ-sensitive proteins by screening a membrane-gene-deficient expression yeast library.** Screening of PCZ resistant yeast strains. Growth of gene-deficient yeast strains on YPDA medium containing DMSO, 0.5 μM PCZ, 1.0 μM PCZ, or 10.0 μM PCZ. Plates were photographed after 3 days of incubation at 30°C.

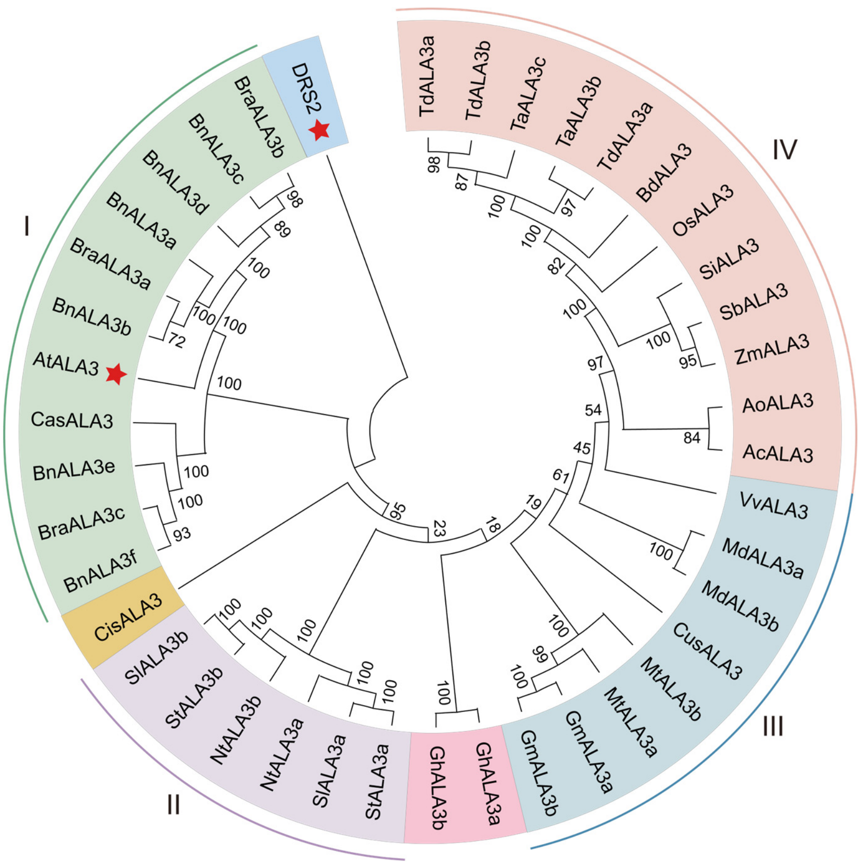


**Fig. S3.**

**Phylogenetic analysis of ALA3 proteins.** Phylogenetic relationships analyzed by neighbor-joining method. Multiple sequence alignment was performed using ClustalW in MEGA 11.0 with default parameters. Bootstrap values from 1,000 replicates are shown at branch nodes.

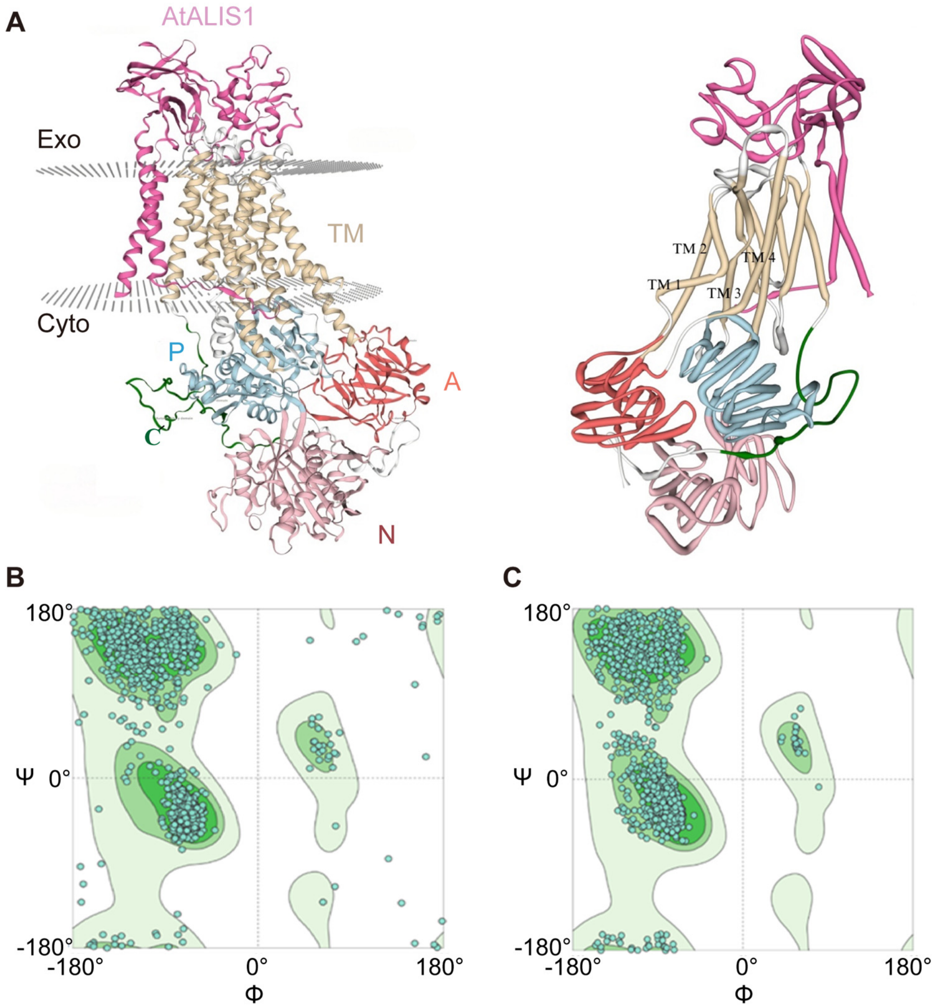


**Fig. S4.**

**Construction and optimization of AtALA3-AtALIS1 protein model. (A)** AtALA3-AtALIS1 protein model. Rose area represents AtALIS1, yellow area represents transmembrane helix TM region, blue area represents P-domain, brick red area represents A-domain, pink area represents N-domain, and green area represents Autoinhibitory C terminus. The plane formed by grey dots is cell membrane region. **(B-C)** Comparison of Ramachandran plots before and after optimization of AtALA3-AtALIS1 protein structure. The figure (B) shows the Ramachandran plot before optimization and the figure (C) shows the Ramachandran plot after optimization. The different regions on the Ramachandran plot reflect the tendency of the amino acid residues to be distributed with respect to the dihedral angles φ and ψ of the amino acid residues' backbone, where the white regions represent the restriction zones, and the degree of green coloring represents the degree of acceptability of the residues.

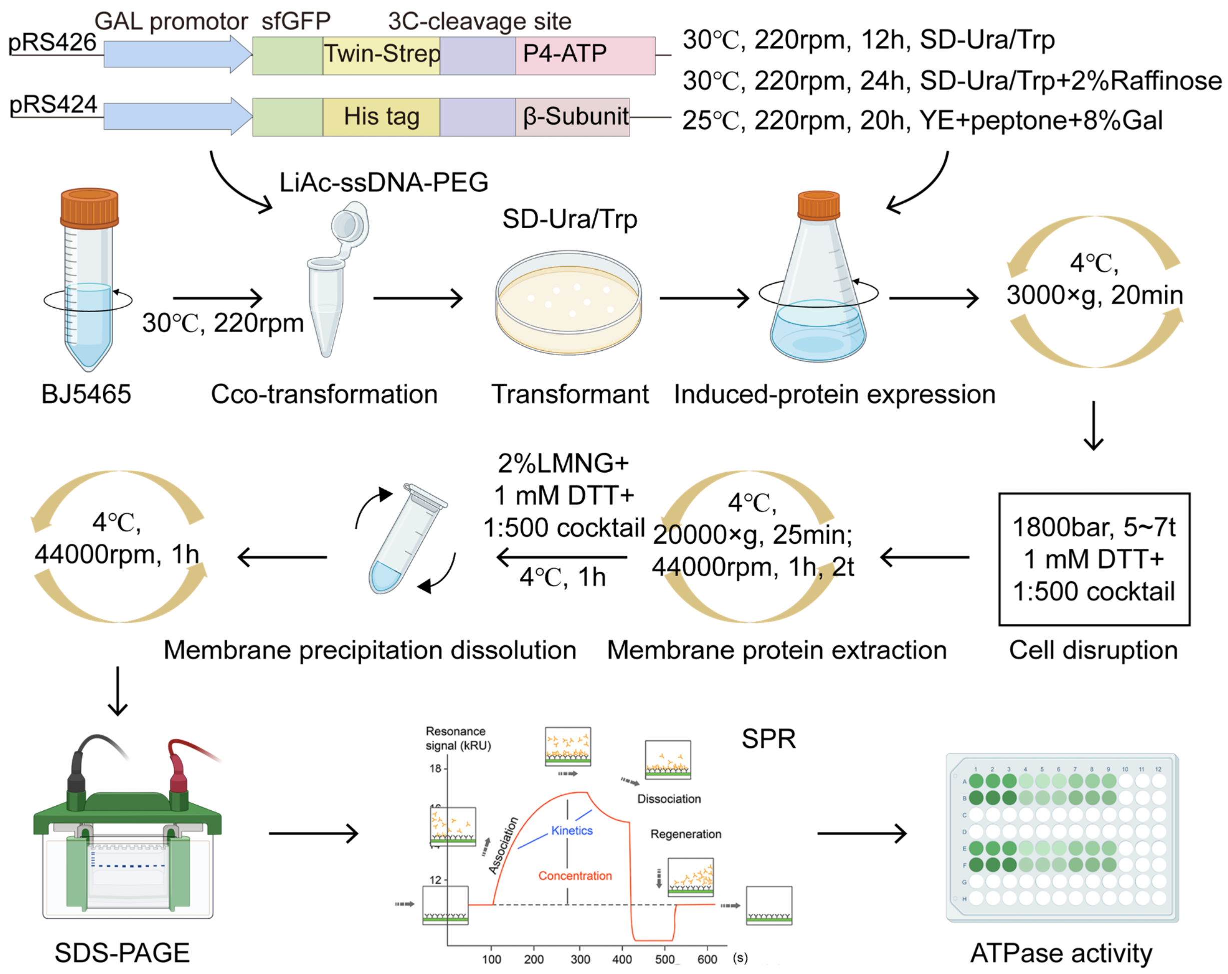


**Fig. S5.**

**Schematic of P4-ATPase protein expression. Protein expression of P4-ATPase.** Schematic of P4-ATPase protein expression and SPR-based affinity measurement. *ALA3* orthologs and their interacting subunits were cloned into pRS426 and pRS424 vectors. Constructs were co-transformed into *Saccharomyces cerevisiae* BJ5465 (a protease-deficient strain with high protein yield and auxotrophic markers trp1, ura3). Transformants were grown in synthetic dropout medium (-Trp/-Ura) to OD600 ≈ 5.0, followed by induction with 8% galactose (25°C, 20 h). Membrane fractions were isolated via differential centrifugation (44,000 rpm, 1 h, 2t) and solubilized in 2% lauryl maltose neopentyl glycol (LMNG). Solubilized proteins were analyzed by SDS-PAGE (Coomassie staining) to confirm expected molecular weights. Protein concentration was determined by BCA assay using BSA as standard. PCZ was immobilized as the stationary phase, and protein was introduced as the mobile phase for surface plasmon resonance (SPR) affinity testing. Data were fitted to a 1:1 Langmuir binding model to calculate ka (association rate), kd (dissociation rate), and KD (equilibrium constant). Using a phosphate assay kit, ATPase activity of the protein complexes under varying conditions was quantified by measuring the absorbance at 620 nm following chromogenic detection of inorganic phosphate released after ATP hydrolysis.

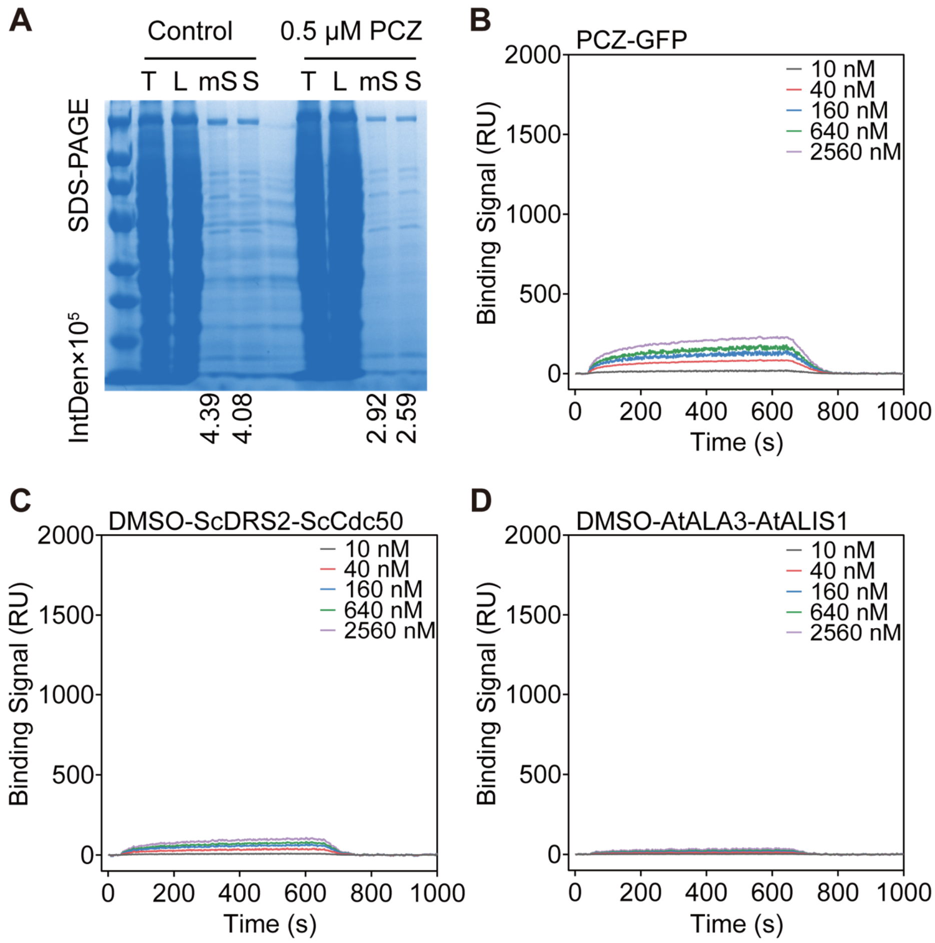


**Fig. S6.**

**SDS-PAGE of AtALA3-AtALIS1 protein expression and SPR affinity binding signals of negative control. (A)** SDS-PAGE of galactose-induced (containing DMSO or 0.5 μM PCZ) AtALA3-AtALIS1 protein expression. T: Total; L: Lysate; mS: membrane Solubilization; S: Supernatant. **(B-D)** SPR affinity binding signals of negative control.

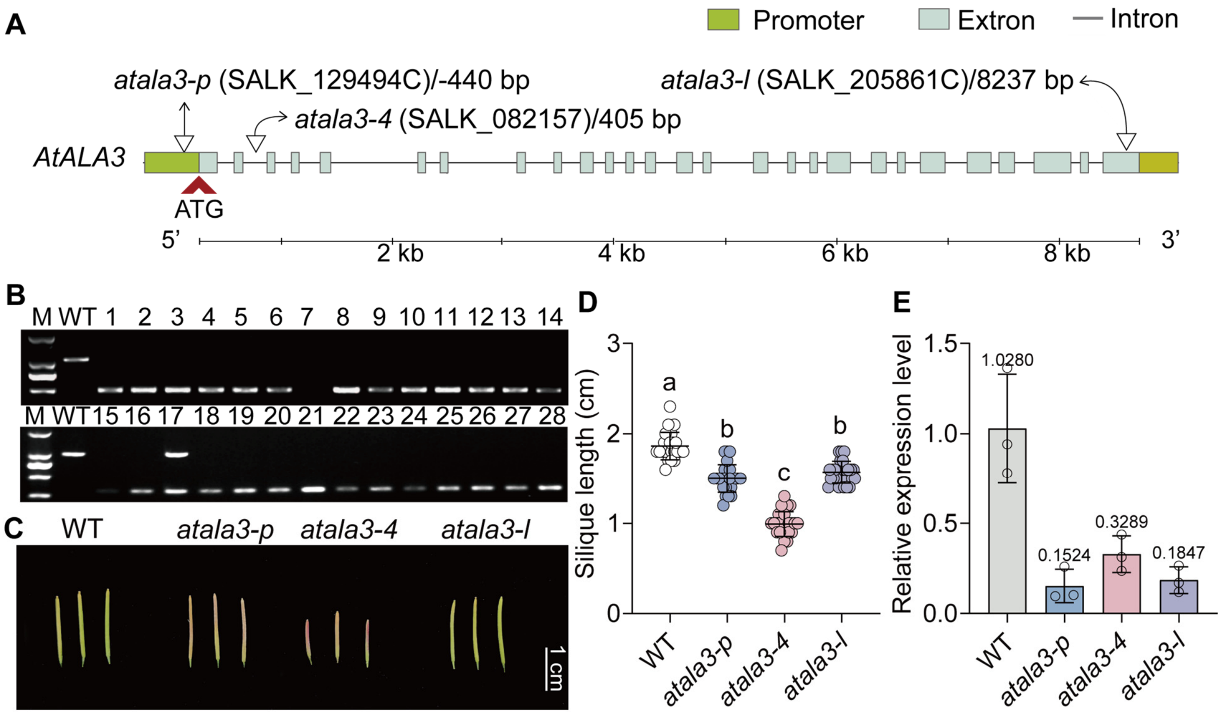


**Fig. S7.**

**Identification and phenotype of *atala3* mutant. (A)** Schematic of *AtALA3* mutant *atala3-p* (SALK_129494C), *atala3-4* (SALK_082157), and *atala3-l* (SALK_205861C) T-DNA insertion sites. "ATG" denotes the translation start site of *AtALA3*. *atala3-p* carries a T-DNA insertion in the promoter region, 440 bp upstream of "ATG". *atala3-4* has an insertion in the second intron, 405 bp downstream of "ATG". *atala3-l* contains an insertion in the last exon, 8,237 bp downstream of "ATG". **(B)** Identification of *atala3-p/l* mutant by triple priming method. **(C)** Silique phenotypes of WT and *atala3-p/4/l* mutant. Scale bar = 1 cm. **(D)** Comparison of pod length of WT and *atala3-1/4/l* mutant. **(E)** Comparison of *AtALA3* relative expression levels in WT and *atala3-p/4/l* mutant (28-day-old). In **D**, the error bar indicates the SD ($n \geq 20$). Letters indicate significant differences between mean values (one-way ANOVA: Tukey, $P < 0.05$). In **E**, the error bar indicates the SD ($n = 3$). Temporal expression profiles normalized to WT.

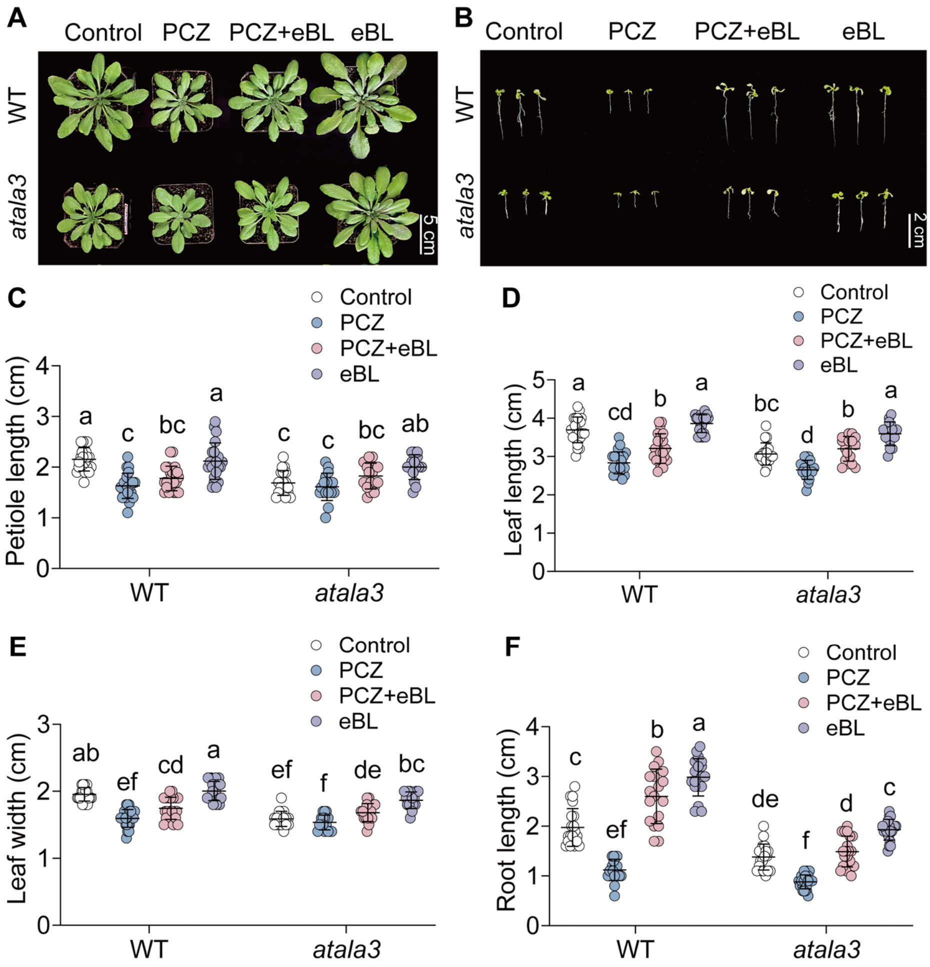


**Fig. S8.**

**Responses of WT and *atala3* to PCZ treatments and eBL complementation. (A)** 5 d phenotypes of DMSO, 50 mg/L PCZ, 50 mg/L PCZ+17 mg/L eBL, and 17 mg/L eBL-treated WT/*atala3* (28-day-old). Scale bar = 5 cm. **(B)** 5 d phenotypes of WT/*atala3* grown on 1/2 MS medium containing DMSO (control), 1 μM PCZ, 1 μM PCZ+0.1 μM eBL, and 0.1 μM eBL. Scale bar = 2 cm. **(C-E)** Comparison of 5 d petiole height (C), leaf length (D), and leaf width (E) of DMSO, 50 mg/L PCZ, 50 mg/L PCZ, PCZ+17 mg/L eBL, and 17 mg/L eBL-treated WT/*atala3*. **(F)** Comparison of 5 d root length of WT/*atala3* grown on 1/2 MS medium containing DMSO (control), 1 μM PCZ, 1 μM PCZ+0.1 μM eBL, and 0.1 μM eBL. In **C-F**, the error bar indicates the SD ($n \geq 15$). Letters indicate significant differences between mean values (one-way ANOVA: Tukey, $P < 0.05$).

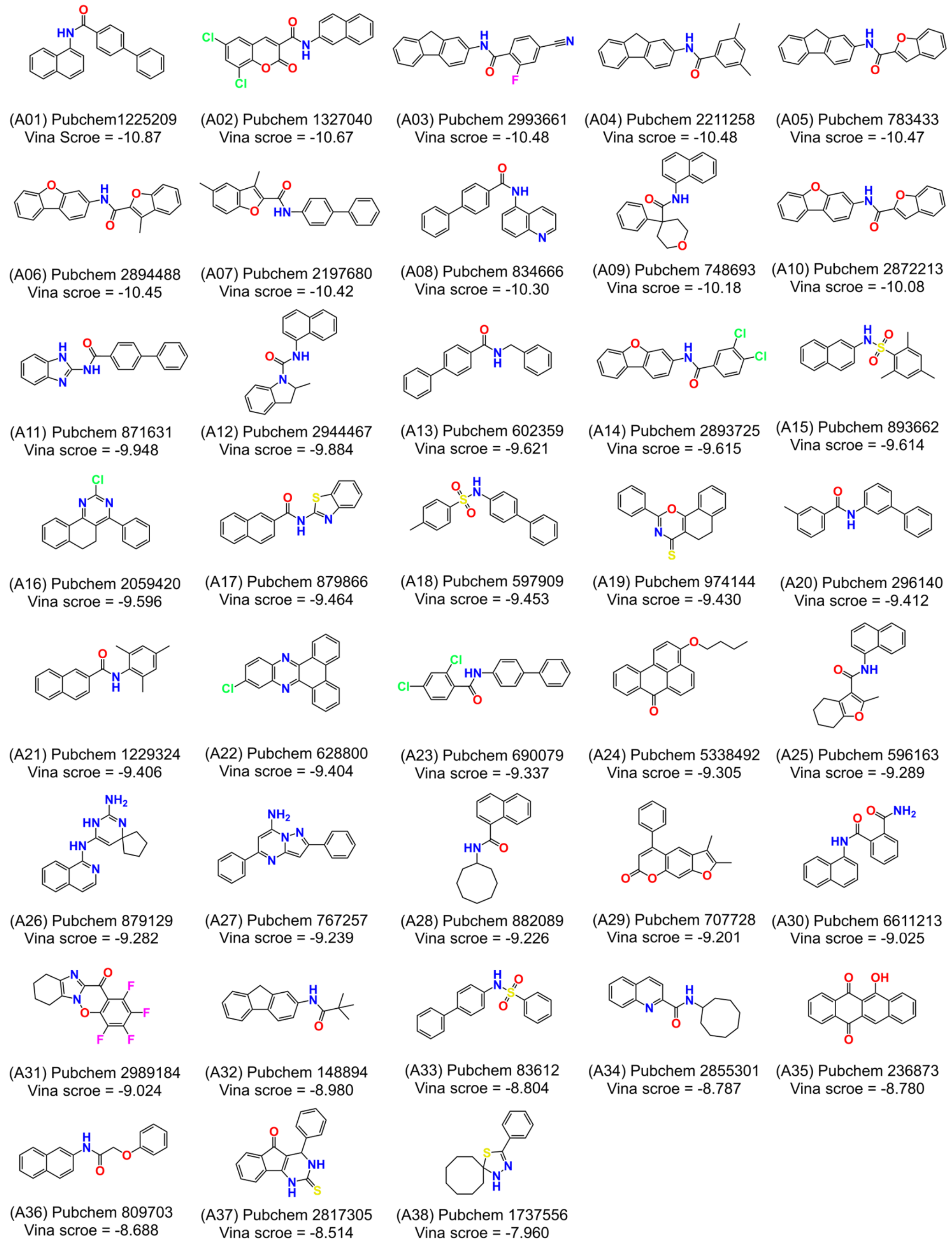


**Fig. S9.**

**Structural formulas and docking scores of 38 compounds obtained by virtual screening targeting the ALA3 active pocket.** A01-A14, A17, A20, A21, A23, A25, A28, A30, A32, A34, and A36 are amide compounds, A15, A18, and A33 are sulfonamide compounds, others are heterocyclic compounds. Structure elements critical for activity have been color-coded: nitrogen atoms(blue), chlorine atoms (green) of the phenyl ring, oxygen atoms (red), fluorine atom (purple), and sulfur atoms (yellow). Structures were drawn using the ChemDraw 22 software and structures were compared to the PubChem database.

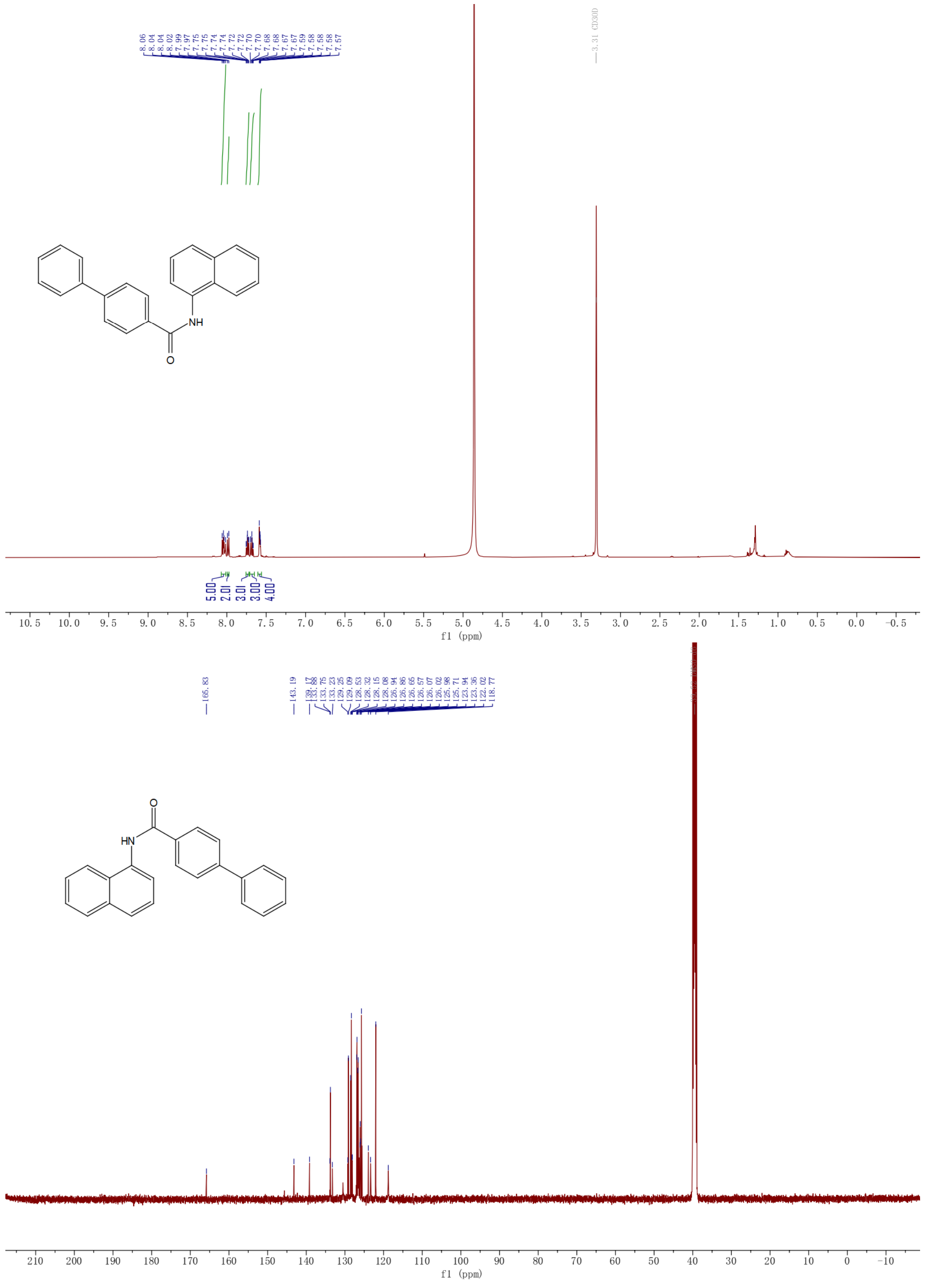


**Fig. S10.**

**Nuclear magnetic resonance (NMR) spectrum of lead compound A01.** $^{1}$H NMR (500 MHz, Methanol-$d_4$) δ 8.04 (dd, $J$ = 12.2, 8.4 Hz, 5H), 7.98 (d, $J$ = 8.4 Hz, 2H), 7.76 – 7.72 (m, 3H), 7.71 – 7.65 (m, 3H), 7.61 – 7.56 (m, 4H). $^{13}$C NMR (126 MHz, DMSO-$d_6$) δ 165.83, 143.19, 139.17, 133.88, 133.75, 133.23, 129.25, 129.09, 128.53, 128.32, 128.15, 128.08, 126.94, 126.86, 126.65, 126.57, 126.07, 126.02, 125.98, 125.71, 123.94, 123.36, 122.02, 118.77.

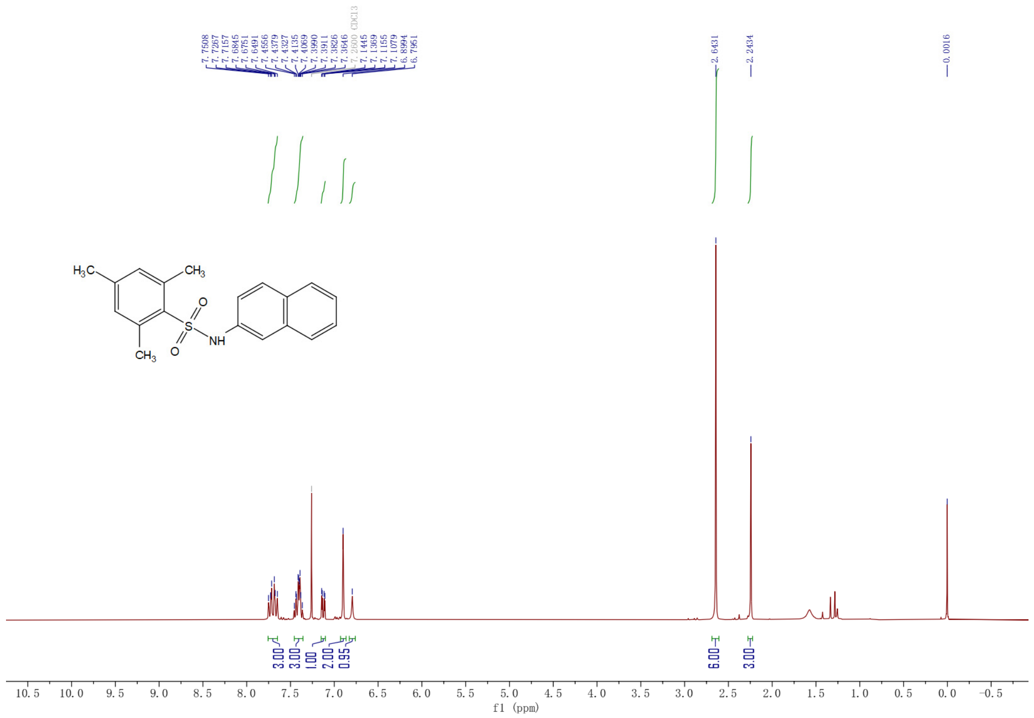


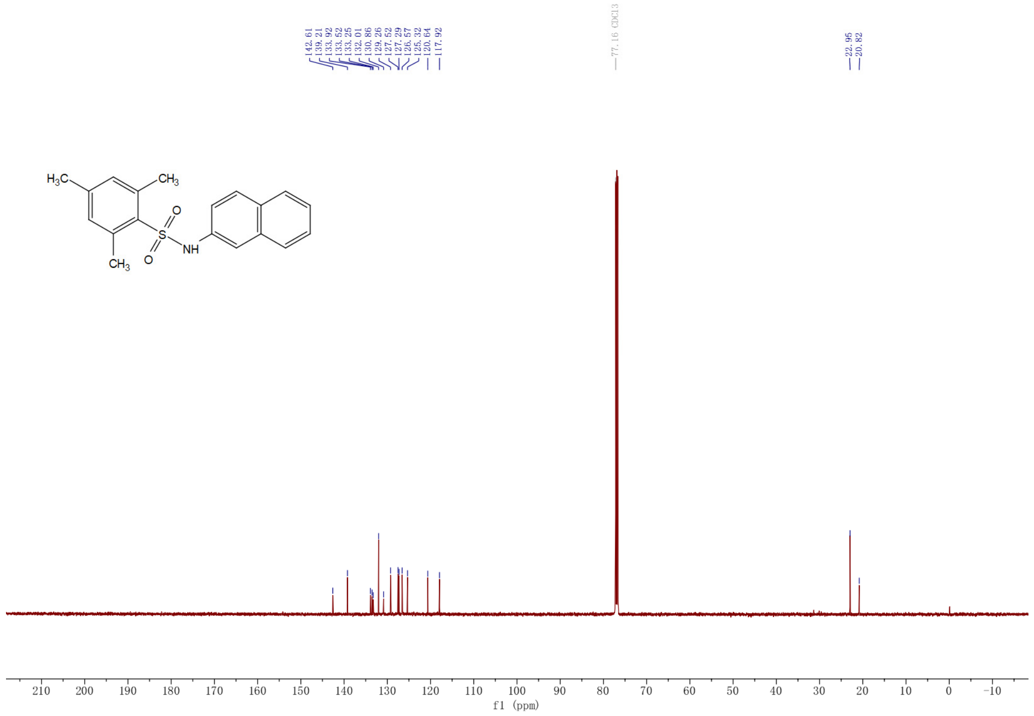


**Fig. S11.**

**NMR spectrum of lead compound A15.** $^{1}$H NMR (300 MHz, Chloroform-*d*) δ 7.75 – 7.65 (m, 3H), 7.46 – 7.36 (m, 3H), 7.13 (dd, *J* = 8.7, 2.3 Hz, 1H), 6.90 (s, 2H), 6.80 (s, 1H), 2.64 (s, 6H), 2.24 (s, 3H). $^{13}$C NMR (126 MHz, Chloroform-*d*) δ 142.61, 139.21, 133.92, 133.52, 133.25, 132.01, 130.86, 129.26, 127.52, 127.29, 126.57, 125.32, 120.64, 117.92, 22.95, 20.82.

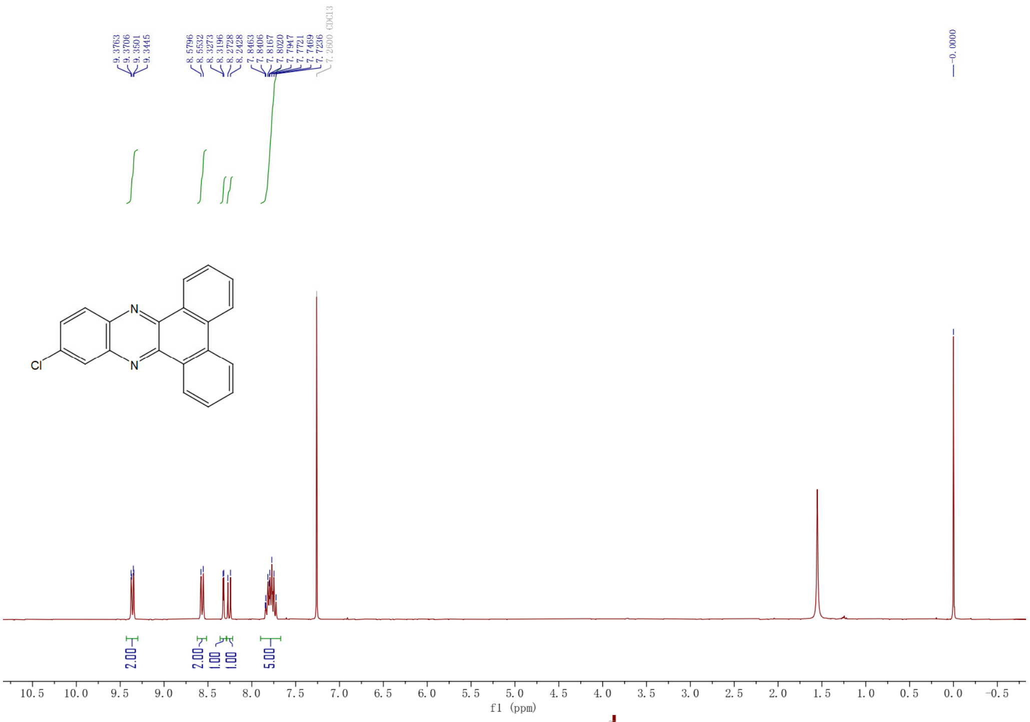


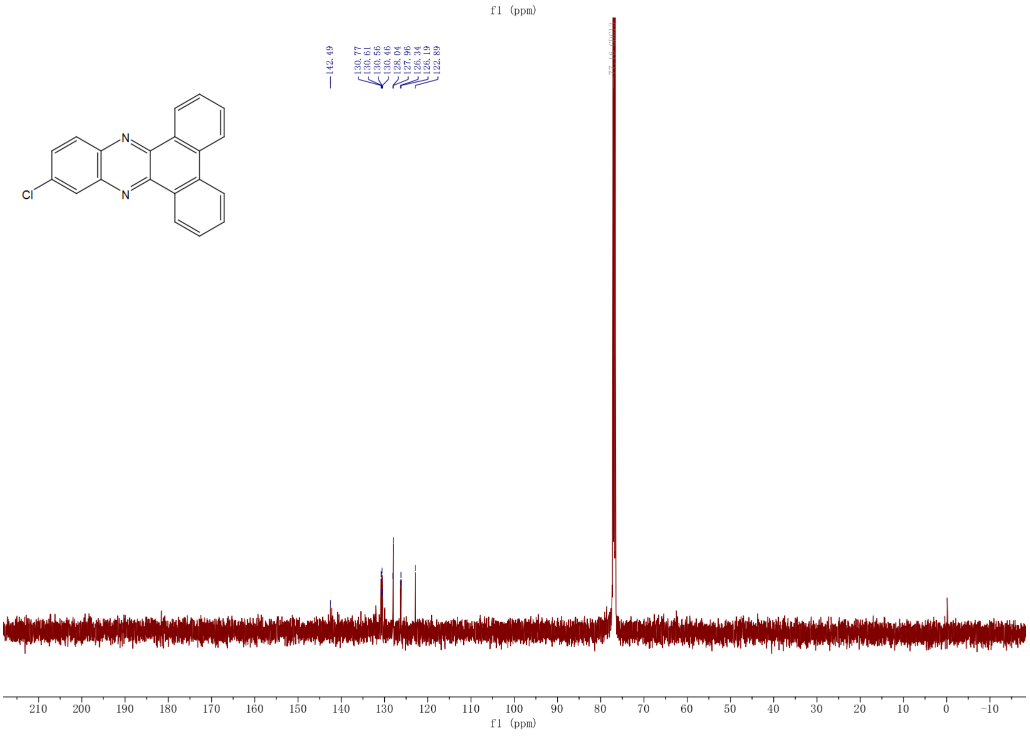


**Fig. S12.**

**NMR spectrum of lead compound A22.** $^1$H NMR (300 MHz, Chloroform-*d*) δ 9.36 (dd, *J* = 7.8, 1.7 Hz, 2H), 8.57 (d, *J* = 7.9 Hz, 2H), 8.32 (d, *J* = 2.3 Hz, 1H), 8.26 (d, *J* = 9.0 Hz, 1H), 7.90 – 7.67 (m, 5H). $^{13}$C NMR (126 MHz, Chloroform-d) δ 130.88, 130.71, 130.67, 130.57, 128.15, 128.06, 126.45, 126.30, 123.00.

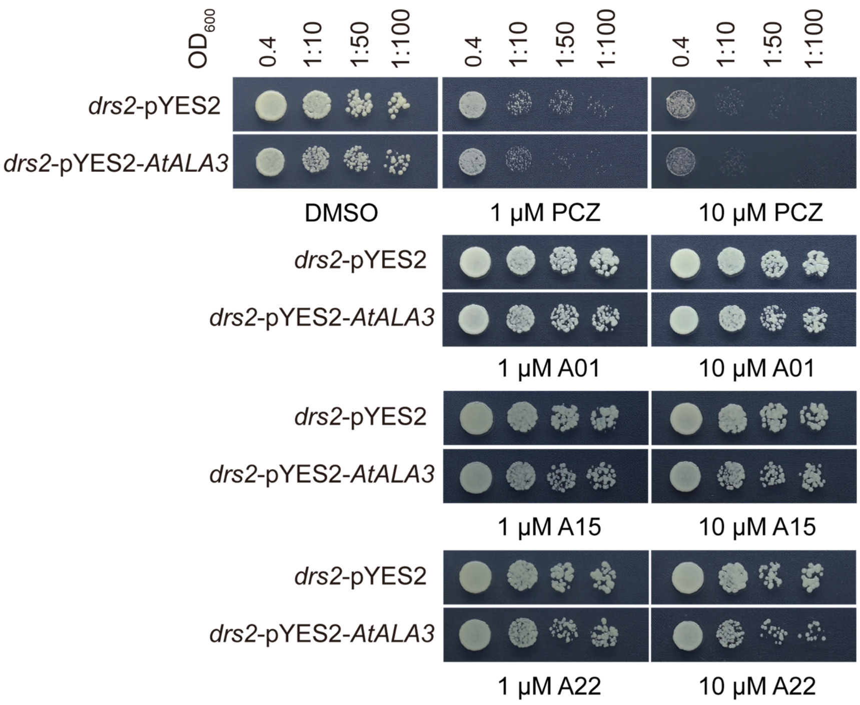


**Fig. S13.**

**Heterologous validation of *AtALA3* sensitivity to three lead compounds in *drs2* yeast strain.** Growth of *drs2* yeast strains supplemented homologous gene *AtALA3* on SD/-His-Leu-Met medium containing DMSO, PCZ, A01, A15, or A22. Plates were photographed after 3 days of incubation at 30°C.

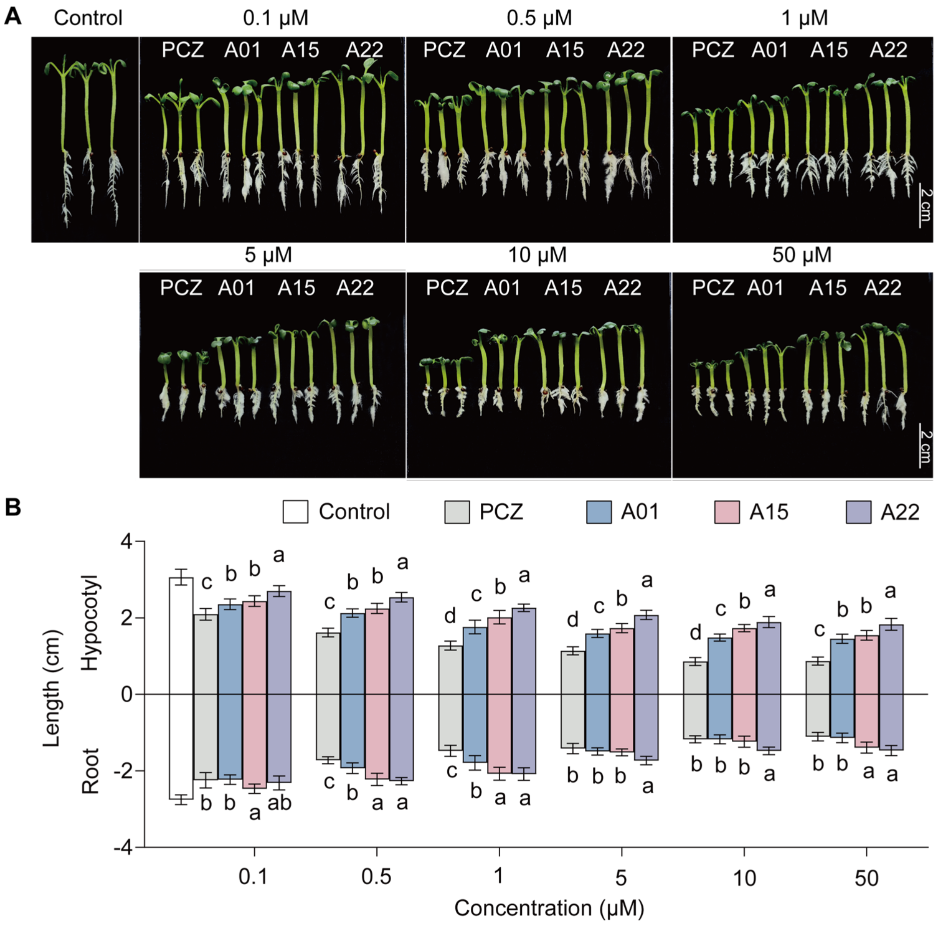


C

| Compound | 0.1 μM | 0.5 μM | 1 μM | 5 μM | 10 μM | 50 μM |
|---|---|---|---|---|---|---|
| | Hypocotyl inhibition rate (%) | | | | | |
| PCZ | 31.67 ± 4.89 | 47.24 ± 3.73 | 58.30 ± 3.81 | 62.82 ± 3.41 | 71.86 ± 3.41 | 71.60 ± 3.40 |
| A01 | 23.13 ± 4.54 | 30.66 ± 3.57 | 42.47 ± 5.89 | 48.00 ± 3.39 | 51.51 ± 2.94 | 52.50 ± 3.90 |
| A15 | 20.36 ± 4.53 | 26.64 ± 4.34 | 34.18 ± 5.79 | 43.48 ± 3.86 | 43.48 ± 3.09 | 49.51 ± 4.13 |
| A22 | 11.82 ± 4.62 | 17.10 ± 4.12 | 26.14 ± 3.14 | 32.17 ± 4.03 | 38.20 ± 4.62 | 40.21 ± 4.62 |
| | Root inhibition rate (%) | | | | | |
| PCZ | 18.44 ± 7.35 | 37.43 ± 3.37 | 46.37 ± 5.17 | 48.61 ± 4.88 | 57.26 ± 3.37 | 59.8 ± 4.00 |
| A01 | 19.00 ± 4.54 | 29.89 ± 5.00 | 34.92 ± 6.87 | 45.81 ± 3.46 | 57.26 ± 4.23 | 58.66 ± 4.58 |
| A15 | 10.34 ± 4.54 | 19.28 ± 5.77 | 24.59 ± 6.14 | 44.98 ± 3.26 | 55.03 ± 5.25 | 49.44 ± 5.23 |
| A22 | 15.93 ± 6.60 | 17.60 ± 3.44 | 24.31 ± 5.90 | 37.15 ± 4.03 | 46.37 ± 3.68 | 46.65 ± 4.78 |

**Fig. S14.**

**Responses of *Brassica rapa* var. *parachinensis* (*B. rapa*) to three lead compounds treatments. (A)** 5 d phenotypes of *B. rapa* grown on 1/2 MS medium containing DMSO (control), PCZ, A01, A15, and A22. Scale bar = 2 cm. **(B)** Comparison of 5 d hypocotyl length and root length of *B.*

*rapa* grown on 1/2MS medium containing DMSO (control), PCZ, A01, A15, and A22. In **B**, the error bar indicates the SD ($n$ = 13). Letters indicate significant differences between mean values (one-way ANOVA: Tukey, $P < 0.05$). **(C)** Statistical analysis of hypocotyl length and root length inhibition rates in *B. rapa* treated with PCZ, A01, A15, or A22, compared to the DMSO control. Data are presented as the mean ± SD.

**Table S1.**

**Detailed information on ALA3s used phylogenetic tree.**

| Species | Gene name | Gene ID |
|---|---|---|
| *Brachypodium distachyon* | *BdALA3* | LOC100846736 |
| *Sorghum bicolor* | *SbALA3* | LOC8064926 |
| *Zea mays* | *ZmALA3* | LOC103643580 |
| *Setaria italica* | *SiALA3* | LOC101777430 |
| *Triticum aestivum* | *TaALA3a* | LOC123042727 |
| | *TaALA3b* | LOC123118639 |
| | *TaALA3c* | LOC123180881 |
| *Triticum dicoccum* | *TdALA3a* | LOC119331482 |
| | *TdALA3b* | LOC119266663 |
| *Asparagus officinalis* | *AoALA3* | LOC109851217 |
| *Oryza sativa* | *OsALA3* | LOC4348609 |
| *Ananas comosus* | *AcALA3* | LOC109718308 |
| *Glycine max* | *GmALA3a* | LOC100815547 |
| | *GmALA3b* | LOC100787473 |
| *Solanum tuberosum* | *StALA3a* | LOC102587636 |
| | *StALA3b* | LOC102585532 |
| *Medicago truncatula* | *MtALA3a* | LOC11408682 |
| | *MtALA3b* | LOC11407613 |
| *Nicotiana tabacum* | *NtALA3a* | LOC104107296 |
| | *NtALA3b* | LOC104101336 |
| *Camelina sativa* | *CasALA3* | LOC104768800 |
| *Vitis vinifera* | *VvALA3* | LOC100264484 |
| *Solanum lycopersicum* | *SlALA3a* | LOC101248674 |
| | *SlALA3b* | LOC101243809 |
| *Malus domestica* | *MdALA3a* | LOC103452351 |
| | *MdALA3b* | LOC103431918 |
| *Cucumis sativus* | *CusALA3* | LOC101207069 |
| *Brassica napus* | *BnALA3a* | LOC106450257 |
| | *BnALA3b* | LOC106376745 |
| | *BnALA3c* | LOC106375547 |
| | *BnALA3d* | LOC106396449 |
| | *BnALA3e* | LOC106346525 |
| | *BnALA3f* | LOC106420125 |
| *Brassica rapa* | *BraALA3a* | LOC103829275 |
| | *BraALA3b* | LOC103862629 |
| | *BraALA3c* | LOC103843300 |
| *Citrus sinensis* | *CisALA3* | LOC102627471 |
| *Gossypium hirsutum* | *GhALA3a* | LOC107959952 |
| | *GhALA3b* | LOC107903682 |
| | *GhALA3c* | LOC107886595 |
| *Saccharomyces cerevisiae* | *DRS2* | YAL026C |

**Table S2.**
**Primers used in this study.**

| Primer name | Primer sequence (5' to 3') |
|---|---|
| **Yeast expression** | |
| pYES2-AtALA3-F | actatagggaatattaagcttATGGTTCGATCGGGTAGTTTTAG |
| pYES2-AtALA3-R | tgatggatatctgcagaattcTTACTTCTTCGGTACCTTTGGC |
| pYES2-357-F | TTTCGGTTTGTATTACTTCTTATTC |
| AtALA3-R | GACATGAAAAGAATATCTGCTGG |
| **RT-qPCR** | |
| Actin-F | CATCAGGAAGGACTTGTACGG |
| Actin-R | GATGGACCTGACTCGTCATAC |
| qRT-AtALA3-F | AGGAGCTTGGACAGGTGGAA |
| qRT-AtALA3-R | TCTTATTGCACCCGTGGACC |
| **Protein expression** | |
| 424-RA-R | TGGACCTTGAAACAAAACTTCCAA |
| 424-RA-F | CTCGAGTCATGTAATTAGTTATGTC |
| pRS424-CDC50-F | caccacttggaagttttgtttcaaggtccaATGGTTTCATTGTTCAAAAGAGG |
| pRS424-CDC50-R | cgtgacataactaattacatgactcgagCTATAAAATTTCCCTCAATGTTGTAT |
| pRS424-AtALIS1-F | aagttttgtttcaaggtccaggcgcgcctATGTCTTCTTCTAACACGCC |
| pRS424-AtALIS1-R | aactaattacatgactcgagggcgcgcctTTAACGACCTCCAGGAATTC |
| CDC50-R | TCGGTTAGCCTCCATTGTGG |
| AtALIS1-R | TGGCTGCCCACCAAAATCATC |
| 426-RA-R | TGGACCTTGAAACAAAACTTCCAACTT |
| 426-RA-F | CTCGAGTCATGTAATTAGTTATGTCACG |
| pRS426-DRS2-F | gaaaagttggaagttttgtttcaaggtccaATGAATGACGACAGAGAAACCC |
| pRS426-DRS2-R | cgtgacataactaattacatgactcgagTCATATATCAAATGAAATATCATCTCT |
| pRS426-AtALA3-F | gaaaagttggaagttttgtttcaaggtccaATGGTTCGATCGGGTAGTTTTAG |
| pRS426-AtALA3-R | cgtgacataactaattacatgactcgagTTACTTCTTCGGTACCTTTGGCC |
| DRS2-R | TGCGTCGCCAACGTTCTTTC |
| **Site-directed mutagenesis** | |
| AtALA3-N1044A-F | GCGCATTCTTTTGATGAGCGCTTCCATTACCAGATGGCAT |
| AtALA3-N1044A-R | ATGCCATCTGGTAATGGAAGCGCTCATCAAAAGAATGCGC |

| Primer name | Primer sequence (5' to 3') |
|---|---|
| **Site-directed mutagenesis** | |
| AtALA3-F1112G-F | TCTTCTAGGCGATTTCATCGGCCAAGGGGTGGAGAGATGG |
| AtALA3-F1112G-R | CCATCTCTCCACCCCTTGGCCGATGAAATCGCCTAGAAGA |
| AtALA3-Q1113A-F | TCTAGGCGATTTCATCTTCGCAGGGGTGGAGAGATGGTTC |
| AtALA3-Q1113A-R | GAACCATCTCTCCACCCCTGCGAAGATGAAATCGCCTAGA |
| AtALA3-Y1124A-F | ATGGTTCTTCCCGTATGATGCTCAGATCGTTCAAGAAATA |
| AtALA3-Y1124A-R | TATTTCTTGAACGATCTGAGCATCATACGGGAAGAACCAT |
| **Genotyping** | |
| AtALA3-1-LP | TCGGAATTCCAAATGACGTAC |
| AtALA3-1-RP | ACAAATTGATTCGATTGCGAG |
| AtALA3-2-LP | TCATCAGGCAAAGTATTTGGC |
| AtALA3-2-RP | AAGTATTGGGTTCCTAACGCC |
| LBa1.3 | ATTTTGCCGATTTCGGAAC |
| **LUC** | |
| 1300-AtALIS1-F | atcctctagagtcgacATGTCTTCTTCTAACACGCCATCTTCT |
| 1300-AtALIS1-R | tacgaacgaaagctctgcagTTAACGACCTCCAGGAATTCTGTTCCAC |
| 1300-AtALA3-CLucF | acgcgtcccggggcggtaccATGGTTCGATCGGGTAGTTTTAGC |
| 1300-AtALA3-CLucR | agctctgcaggtcgacTTACTTCTTCGGTACCTTTGGCCG |
| CLuc-F | CCAGTCAAGTAACAACCGCGA |
| 1300-AtCYP51G1-NLucF | cgggggacgagctcggtaccATGGAATTGGATTCGGAGAACAAATTGT |
| 1300-AtCYP51G1-NLucR | acgagatctggtcgacAGAAAGCTGGCGCCTCT |
| NLuc-R | CCTTATGCAGTTGCTCTCCAG |
| AtCYP51G1-F | TCAAAGACCCCGACACCTACG |
| **BiFC** | |
| mycYFPN-AtALA3-F | tctatatcatggccggtaccATGGTTCGATCGGGTAGTTTTAGC |
| mycYFPN-AtALA3-R | agctctgcaggtcgacTTACTTCTTCGGTACCTTTGGCCG |
| YFPN-F | AACTACAAGACCCGCGCCG |
| HAYFPC-AtCYP51G1-F | acgagctgtacaagggtaccATGGAATTGGATTCGGAGAACAAATTGT |
| HAYFPC-AtCYP51G1-R | AAGCTctgcaggtcgacTTAAGAAAGCTGGCGCCTCTTGTAAC |
| YFPC-F | GACAACCACTACCTGAGCTAC |

| Primer name | Primer sequence (5' to 3') |
|---|---|
| **Y2H** | |
| pBT3N-AtALA3-F | atatcgaattcctgcagggccattacggccATGGTTCGATCGGGTAGTTTTAGC |
| pBT3N-AtALA3-R | ttagctacttaccatggggccgaggcggccCTTCTTCGGTACCTTTGGCCGTGAT |
| pBT3-N-F | CAGAAGGAGTCCACCTTACAT |
| pBT3-N-R | GGGACCTAGACCTTCAGGTTG |
| pPR3-C-AtCYP51G1-F | gtatcaacgcagagtggccattacggccATGGAATTGGATTCGGAGAACAAATTGT |
| pPR3-C-AtCYP51G1-R | atcgaattctcgagaggccgaggcggcAGAAAGCTGGCGCCTCTTGTAACGC |
| pPR3-C-F | CGGCCTTCCTTCCAGTTACTTG |
| pPR3-C-R | CGTTGTCGATGGTATCGGAAGAT |
| **CO-IP** | |
| 1307myc-AtALA3-F | gtatctagaactagtggatccATGGTTCGATCGGGTAGTTTTAG |
| 1307myc-AtALA3-R | gggcccccctcgaggtcgacTTACTTCTTCGGTACCTTTGGCCG |
| 1300-AtCYP51G1-3flag-F | tggagagaacacgggggacgagctcATGGAATTGGATTCGGAGAACAAATTGTTG |
| 1300-AtCYP51G1-3flag-R | ttgtagtccatgtcgactctagaggatccAGAAAGCTGGCGCCTCTTGTAACGC |
| **Gene silencing** | |
| pre-miRNA-F | ggactctagaggatccGGGTGAGAATCTCCATGT |
| pre-miRNA-R | gatcggggaaattcgagctcGGGTGAAGAGCTCATGT |
| G418-F | CGGCTATGACTGGGCACAACAGACAAT |
| G418-R | CTCGGCAGGAGCAAGGTGAGATGAC |
| qRT-BraUBC10-F | GGGTCCTACAGACAGTCCTTAC |
| qRT-BraUBC10-R | ATGGAACACCTTCGTCCTAAA |
| qRT-BraALA3a-F | ACACCACGTGATAGAAATGAAAATG |
| qRT-BraALA3a-R | CGCCTAGAAGAGAAACAATGGG |
| qRT-BraALA3b-F | CGATTTCATCTACCAAGGGGTT |
| qRT-BraALA3b-R | TGCTTTGAGAGCTCCCGT |
| qRT-BraALA3c-F | CGCCACGTGATAGAAACGAGA |
| qRT-BraALA3c-R | GAAGAACCACCTCTCCACCC |

**Table S3.**

**Site-directed mutagenesis information of AtALA3 heterologous expression in yeast.**

| | Codon position | Codon | Mutagenesis genesis Codon | Amino acid position | Amino acid | Mutagenesis Amino acid | Polarity | Muta-genesis Polarity |
|---|---|---|---|---|---|---|---|---|
| N1044A | 3132 | AAT | GCT | 1044 | ASN | ALA | p | np |
| F1112G | 3336 | TTC | GGC | 1112 | PHE | GLY | np | p |
| Q1113A | 3339 | CAA | GCA | 1113 | GLN | ALA | p | np |
| Y1124A | 3372 | TAT | GCT | 1124 | TYR | ALA | p | np |

"p" indicates amino acids containing polar and neutral R groups, "np" indicates amino acids containing non-polar and hydrophobic R groups.

**Table S4.**
**SPR binding affinities between AtALA3 protein and PCZ, A01, A15, A22.**

| Compound | Protein | Avg ka (1/Ms) | Avg kd (1/s) | **Avg KD (M)** | Int.Intensity Level | ABS (tr_KD) |
|---|---|---|---|---|---|---|
| PCZ | AtALA3 | 2.06E+05 | 4.21E-03 | 2.04E-08 | Strong | 25.5450 |
| A01 | AtALA3 | 2.06E+05 | 1.05E-03 | 5.11E-09 | Strong | 27.5450 |
| A15 | AtALA3 | 3.03E+05 | 4.07E-03 | 1.34E-08 | Strong | 26.1496 |
| A22 | AtALA3 | 4.88E+04 | 5.74E-03 | 1.18E-07 | Strong | 23.0182 |
| PCZ | GFP | 4.32E+00 | 3.73E-01 | 8.62E-02 | VW/None | 3.5359 |
| DMSO | AtALA3 | 1.24E+00 | 7.19E-01 | 5.78E-01 | VW/None | 0.7903 |

SPR affinity coefficients of AtALA3-AtALIS1 and PCZ, A01, A15, A22. Avg Ka (1/Ms): The average association rate constant (Ka) represents the ratio of complex formation per unit time relative to initial reactant concentrations, where higher values indicate faster molecular binding. Avg Kd (1/s): The mean dissociation rate constant (Kd) reflects the proportion of complex dissociation per unit time, with elevated values corresponding to faster complex disintegration. Avg KD (M): The equilibrium dissociation constant (KD = Kd/Ka) quantifies binding affinity at dynamic equilibrium. Lower KD values signify stronger intermolecular interactions. Interaction Intensity Level: Determination of affinity. KD range: $10^{-13}$-$10^{-5}$ M = strong; $10^{-5}$-$10^{-3}$ M = moderate; $10^{-3}$-$2\times10^{-2}$ M = weak; >$2\times10^{-2}$ M = negligible. ABS (tr_KD): Absolute affinity coefficient. ABS (tr_KD) = ABS ($\log_2$KD), where higher numerical values indicate enhanced binding affinity. Reported as mean ± SD from four technical replicates.

**Method S1.**

**Protein 3D modelling of AtALA3-AtALIS.**

Previous studies have established ALA3 as a P4-ATPase phospholipid flippase characterized by ten transmembrane domains, and P4-ATPases interact with a *β*-subunit to facilitate their function (*17, 18*). Based on reported studies, the final localization and lipid substrate specificity of P4-ATPases are independent of the properties of their cognate ALIS *β*-subunits that mediate these interactions (*27*), we selected *A. thaliana* AtALIS1 as the *β*-subunits for modeling. A homology model of the AtALA3-AtALIS1 complex was generated based on the structure of the DRS2-Cdc50p template. The conformational structures of DRS2-Cdc50p protein were obtained from Protein Data Bank (PDBe, https://www.ebi.ac.uk/pdbe/) (*20*). The 3D structures of the ligand molecules propiconazole (PCZ), and phosphatidylinositol 4-phosphate (PI4P) were downloaded from the PubChem database (https://pubchem.ncbi.nlm.nih.gov/). High-resolution crystal structures have not been reported for AtALA3-AtALIS1 double-stranded proteins, and the artificial intelligence model ColabFold (https://colabfold.mmseqs.com/) was used for protein 3D structure modelling (*56*). The amino acid structures were obtained from the UniProt database (https://www.uniprot.org/), and were hosted on the Google Colaboratory project hosting platform using ColabFold, the structure of the polypeptide chain complexes was predicted by multichain mode, with other parameters defaulted. After submitting the task, ColabFold was trained using transformer model and outputed a predicted structure file (PDB format). The predicted model was reviewed by PyMOL visualisation software. The complex model was evaluated and optimised using the methods of Ramachandran (*57*).

A structural model of the AtALA3-AtALIS1 complex was generated using this approach (fig. S4A). Assembly of AtALIS1 subunit with AtALA3 protein, the constructed AtALA3 AtALIS1 complex model contains 10 transmembrane structures α-Spiral (TM1-10), where TM1-6 and TM7-10 aggregate to form 2 spiral bundles, connected by a larger cytoplasmic ring in the middle, forming the main hydrophobic core of the entire complex. The AtALIS1 subunit contains 2 transmembrane α-helices, *β*-folding and irregular curling of the extracellular region, forming a "cap" structure (fig. S4A). This structural feature is consistent with typical P-type ATPase family members (*58*). Ramachandran plot results show that most of the residues of the pre-optimised structure have their main chain dihedral angles distributed in the acceptable region, while some residues fall into the restricted region (fig. S4B), indicating that there is still room for optimisation of the structure. After the optimisation, no residues were found in the restricted region and the overall concentration of the main chain dihedral angle was towards the most acceptable region (fig. S4C), indicating that the conformational rationality of the structure has been significantly improved, and this structure was used for molecular docking of subsequent protein models.

**Method S2.**

**Molecular docking of AtALA3-AtALIS with PCZ.**

Water molecules and other small-molecule ligands in the protein structure were removed using PyMOL to check the protein atom type and structural integrity, and then the protein molecules were added with polar hydrogen atoms and Gaussian charges and the ligand small molecules were set up with torsionally reversible bonding positions using AutoDock Tools to obtain a standard PDBQT file. Based on previous reports (*20, 59*), the parameters of the docking box were determined (box dimensions: 30 Å × 30 Å × 30 Å, lattice length: 0.375 Å). Molecular docking was performed using Autodock Vina (*60*), vina.exe was used for docking and vina_split.exe was used to split the docking results, exhaustiveness = 100 was setted for more accurate results. The algorithm of Autodock Vina was set as lazy scoring function and genetic algorithm when executing docking to find the best binding conformation and affinity between ligand and receptor in the set search space. The ligand-receptor interactions in each binding mode, including hydrogen bonding, hydrophobic forces, and van der Waals forces between the ligand and active site residues, were analysed by PyMOL and UCSF ChimeraX.

**Method S3.**

**Molecular dynamics simulation (MD).**

50 ns all-atom molecular dynamics simulations of the complex model were performed using the GROMACS software package (*61*). The structure files of proteins and small molecule ligands were uploaded through the Protein/Ligand Complex module of the CHARMM-GUI online server (https://charmm-gui.org/), and the structures were processed for centre of mass alignment, residue error correction, and missing atom construction, and the system was run through the process of ionisation (0.15 mol/L NaCl) and solvation (TIP3P water model) to construct topology files . Energy minimisation files were generated using the gmx grompp tool of GROMACS, with the maximum number of steps set to 50,000, and conjugate gradient energy minimisation was carried out via the gmx mdrun command with a time step of 1 fs. The NPT equilibrium condition was constant temperature (T = 300 K) and constant pressure (P = 1 bar), and was performed using a V-rescale temperature coupling and a Parrinello-Rahman pressure coupling, with a simulation duration of 100 picoseconds. Molecular dynamics simulations of final conformation production based on NPT equilibrium were performed with gmx grompp to generate input files for 50 ns molecular dynamics production runs. The final molecular dynamics trajectories were used to analyse the structure, conformational dynamics and interaction characteristics of protein-small molecule complexes. The stability of conformations was assessed by Root Mean Square Deviation (RMSD), Root Mean Square Fluctuation (RMSF), andRadius gyration (Rg).

**Method S4.**

**Histological section.**

PCZ-treated *A. thaliana* samples were fixed overnight at 4°C in fixative (2.5% Glutaric dialdehyde, 0.1 M Phosphate Buffer [pH 7.2], 0.1% Tween-20 and 4% Paraformaldehyde), followed by graded ethanol dehydration. The samples were treated with a mixture of acetone and embedding agent with V/V = 3/1, 2/1, 1/1, 1/2, 1/2 and 1/3 for 2 h. Finally, the samples were treated twice with pure embedding agent for 24 h each time. The samples were embedded by Eponate 12™–Araldite embedding Kit with DMP-30 (TED PELLA, INC). The samples were polymerized at 60°C for 16-24 h until the samples cooled to certain hardness. The samples were trimmed to the appropriate size and section, embedded plant tissues were sectioned by the *Leica RM2235* manual microtome in 5-10 μm. The toluidine blue-stained sections were observed under Laser capture micro-dissection (LMD) (*25*). Cell length and width were quantified from stained samples using Image-Pro Plus 6.0 software.

**Method S5.**

**Knockdown of *BraALA3a/b* in *B. rapa*.**

Artificial microRNA (amiRNA) sequences targeting *BraALA3a/b* were designed using WMD3 (http://wmd3.weigelworld.org), validated for specificity via psRNATarget (https://www.zhaolab.org/psRNATarget/), and assessed for secondary structure using the UNAFold Web Server (https://www.unafold.org/). Synthesized fragments (Tsingke Biotechnology Co., Ltd.) were cloned downstream of the CaMV 35S promoter in pBI121 via homologous recombination. The construct was transformed into Agrobacterium tumefaciens GV3101, cultured in LB medium (10 g/L tryptone, 5 g/L yeast extract, 10 g/L NaCl) with antibiotics (50 μg/mL Kan, 50 μg/mL streptomycin [Str], 20 μg/mL rifampicin [Rif]). Cells were harvested by centrifugation and resuspended in infiltration buffer (5% sucrose, 0.02% Silwet L-77, $OD_{600}$ = 1.5). *B. rapa* 'Youqing Sijiu' was transformed via vacuum-infiltration floral dip (-0.09 MPa), with 6-8 manual pollinations using non-transformed pollen (*34*, *35*). After removing secondary bolts, T1 seeds were harvested at 8-10 weeks. Transgenic plants were screened by PCR and validated by qRT-PCR.
